# A simplified protocol for the generation of apical-out human bladder organoids in suspension using a rotating bioreactor

**DOI:** 10.64898/2026.11.23.768490

**Authors:** Valentin Borgeat, Vishwa J. Vasani, Hugo Sallet, Lorenz Leitner, John D. McKinney, Kathrin Tomasek

## Abstract

Bladder organoids are commonly generated in extracellular matrix domes, where apical surfaces line an enclosed central lumen, limiting experimental accessibility. Suspension-based systems have been explored to generate polarity-reversed organoids, but current models are often derived from cancer cells, remain incompletely characterized or rely on highly specialized scaffolds. Here, we describe a rapid and scalable approach to generate apical-out human bladder organoids (hBOs) from primary bladder epithelial cells using a rotating bioreactor system with minimal extracellular matrix scaffolding. Organoids form within 8 days and exhibit an apical-out phenotype with outward-facing umbrella-like cells. These apical-out hBOs tolerate 100% pooled human urine for up to 4 hours and can be infected with uropathogenic *E. coli*, recapitulating hallmarks of infection such as the formation of intracellular bacteria communities. Overall, the apical-out configuration enables direct access to the umbrella cell layer, thereby simplifying experimental manipulation, including infection and treatment assays under physiological conditions.

## Introduction

The bladder epithelium forms a highly specialized, stratified barrier composed of basal, intermediate, and terminally differentiated-umbrella cells, which together maintain tissue integrity and mediate interactions with the urinary environment. Different *in vitro* bladder models, including transwell systems, organoids and organoid-on-a-chip systems, have been developed to recapitulate key aspects of urothelial stratification and differentiation, providing a range of physiologically relevant platforms that reduces reliance on animal models (Jafari and Rohn, 2023; Paduthol et al., 2026; Sharma et al., 2021). These systems have been successfully applied to study bladder cancer and tumor physiology, and host-pathogen interactions, particularly in the context of urinary tract infections (UTIs), which represent one of the most common bacterial infections and are associated with high recurrence rates and increasing antibiotic resistance (Jafari and Rohn, 2022; Vasyutin et al., 2019).

Bladder organoids can be generated using different culture configurations, including extracellular matrix dome cultures (Sharma et al., 2021), as well as suspension-based approaches such as rotation systems (Smith et al., 2006), hanging drop or low-adherent conditions (Amaral et al., 2017; Kim et al., 2020; Vasyutin et al., 2019). In extracellular matrix domes, epithelial cells with stem cell-like properties are embedded and self-organize into 3D structures with a central lumen and an apical-in orientation (Kim et al., 2020; Sharma et al., 2021). In this configuration, terminally differentiated umbrella cells line the luminal surface, thereby restricting direct access to the apical membrane. Consequently, experimental approaches targeting the apical surface, such as drug treatment or infection assays with uropathogenic bacteria, often rely on technically demanding microinjection procedures (Kim et al., 2020; Sharma et al., 2021). In addition, dome-based cultures require substantial amounts of extracellular matrix and media and are limited in scalability, which can constrain their use in high-throughput applications.

Suspension-based culture systems, including rotation cultures, hanging drop and low-adherent environments, offer an alternative that can improve accessibility, scalability and experimental flexibility. Several bladder cancer organoid models generated under such conditions form within short time frames (up to one week), but remain incompletely characterized with respect to urothelial stratification and differentiation (Amaral et al., 2017; Fullerton et al., 2005; Xiao et al., 2005). Notably, two studies have described more extensively characterized systems in which bladder organoids adopt an apical-out phenotype (Ringuette Goulet et al., 2017; Smith et al., 2006). In these models, the apical surface, corresponding to the umbrella cell layer, is directly exposed to the surrounding environment, thereby facilitating functional assays and enabling more physiologically relevant experimental conditions. However, these approaches rely on cancer cell lines and specialized scaffolds, such as a mix of fibroblast- and endothelium-derived air-liquid interfaces or porcine small intestine extracellular matrix, which may limit their broader applicability (Ringuette Goulet et al., 2017; Smith et al., 2006).

Here, we describe a rapid and scalable approach to generate apical-out human bladder organoids (hBOs) from primary bladder epithelial cells using a rotating bioreactor system (CERO 3D, Omni Life Sciences). In this configuration, cells self-assemble in suspension in the presence of low concentrations of matrix, forming organoids with outward-facing umbrella-like cells within 8 days. Compared to conventional dome-based cultures, this approach reduces handling complexity and resource consumption while increasing organoid yield and shortening culture time. Additionally, compared to other suspension-based systems, this approach relies on minimal extracellular matrix support. Most importantly, the apical-out organization provides direct access to the umbrella cell layer, enabling simplified infection and treatment assays of bladder organoids grown from primary cells without the need for microinjection. Together, this system provides a practical and adaptable platform for studying bladder epithelial biology and host-pathogen interactions under controlled conditions. As a proof of concept, we show that these hBOs can survive in 100% pooled human urine over 4 hours and can be used as a model of uropathogenic *E. coli* (UPEC) bladder infections.

## Material and Equipment

### 1.1 Reagents

- Complete Epithelial Cell Medium /w Kit – 500 ml (Cell Biologics, #H6621)
- Fetal Bovine Serum (FBS) (Thermo Fisher, #A5256701)
- MycoZap™ Mycoplasma Elimination Reagent (Lonza, #LT07-918)
- MycoAlert® Mycoplasma Detection Kit (Lonza, #LT07-318)
- DAPI solution (Becton Dickinson, #564907)
- Cultrex Reduced Growth Factor Basement Membrane Extract, Type 2, Pathclear (R&D Systems, #3533-010-02)
- 3dGRO® L-WRN Conditioned Media Supplement (L-WRN) (Sigma-Aldrich, #SCM105)
- TryPLE (Thermo Fisher, #12605010)
- DMEM/F-12, GlutaMAX™ supplement (Thermo Fisher, #31331093)
- B-27™ Supplement (50X), serum free (Thermo Fisher, #17504044)
- Human FGF-7 (KGF), Recombinant Protein, PeproTech® (Thermo Fisher, #100-19-100UG)
- Human FGF-10 Recombinant Protein, PeproTech® (Thermo Fisher, #100-26-100UG)
- A 83-01 (Stemcell Technologies, #100-0245)
- Y-27632 dihydrochloride (Bio-Techne, #1254/50)
- N-Acetyl-L-cysteine (Sigma-Aldrich, #A9165-25G)
- Nicotinamide (Sigma-Aldrich, #N0636-500G)
- PeproGMP® Human VEGF-165 Recombinant Protein, PeproTech® (Thermo Fisher, #GMP100-20-50UG)
- Trypan Blue Stain (Thermo Fisher, #T10282)
- Human Urine Gender Pooled 0.2 μm Filtered Viral Negative (BioIVT, #HUMANURINE-011277)
- Dimethyl Sulfoxide (DMSO) (Chemie Brunschwig, #BP231-100)
- Pierce 16% Formaldehyde (PFA) (w/v), Methanol-free (Thermo Fisher, #28908)
- Glutaraldehyde solution 50 wt. % in H_2_O (Sigma-Aldrich, #340855)
- 0.1% Gelatin-Based Coating Solution (ready-to-use) (Cell Biologics, #6950-100)
- Primocin (InvivoGen, #ant-pm-1)
- Bovine Serum Albumin (BSA), Fraction V (Sigma-Aldrich, #12659-500GM)
- Phosphate buffered saline (PBS), pH 7.4 (Thermo Fisher, #10010-056)
- Phosphate buffer 0.1 M, pH 7.4 (PB buffer; Sigma-Aldrich #P5244)
- HistoGel™ (epredia, #HG-4000-012)
- Wheat Germ Agglutinin (WGA) Alexa Fluor 555 (Thermo Fisher, #W11261)
- DRAQ7 (Thermo Fisher, #D15106)
- NucBlueTM Live ReadyProbes™ Reagent (Hoechst 33342) (Life Technologies, #R37605)
- Sodium chloride (Sigma-Aldrich, #S7653-250G)
- Potassium chloride (Sigma-Aldrich, #60128-250G-F)
- Calcium chloride (CaCl_2_) (Sigma-Aldrich, #C4901-500G)
- Magnesium chloride (Sigma-Aldrich, #M2670-100G)
- HEPES (Thermo Fisher, #15630056)
- RNAeasy Plus Mini Kit (QIAGEN, #74134)
- SuperScript®IV First-Strand Synthesis System with random hexamers (Life Technologies, #18090050)
- dNTP (Thermo Fisher, #18427-013)
- RNaseOUT™ Recombinant Ribonuclease Inhibitor (Thermo Fisher, #10777-019)
- DEPC-treated water (Thermo Fisher, #10813-012)
- SYBRGreen PCR Master Mix (Life Technologies, #4364344)
- ProLong™ Gold Antifade Mountant (Invitrogen™, #P36930)
- Osmium tetroxide (EMS, Hatfield, PA, #19110)
- Potassium hexacyanoferrate (II) trihydrate (Sigma-Aldrich, #60279-250G)
- Agarose, low gelling temperature (Sigma-Aldrich, #A9414-50G)
- Acetone, histological grade, ≥99.5% (Sigma-Aldrich, #534064-500ML)
- Acetate uranyl dihydrate (Sigma-Aldrich, #179337-500ML)
- Sodium citrate dihydrate (Sigma-Aldrich, #S1804-500G)
- Lead (II) nitrate (Merck, #A675698 733)
- Sodium hydroxide, pellets, reagent grade, ACS, ISO, Raeg. Ph Eur, 1Kg (Scharlau, #SO04251000)
- Embed 812 Resin (Electron Microscopy Sciences, #14900)
- Dodenyl Succinic Anhydride, DDSA (Electron Microscopy Sciences, #13710)
- Methyl-5-Norbornene-2,3-Dicarboxylic Anhydride, NMA (Electron Microscopy Sciences, #19000)
- Benzyldimethylamine, BDMA (Electron Microscopy Sciences, #11400)

### 1.2 Biological materials

- Bacterial strain: TS85 (CFT073, att_*HK022*_:: *σ*_70_-*sfGFP*) (Simonet et al., 2022)
- Human cell line: Human Primary Bladder Epithelial Cells (Cell Biologics, #H-6215, Lot: F051616)
- Bladder tissue specimens: Obtained during clinically indicated transurethral procedures independent of study participation at Balgrist University Hospital, either by cold-cup biopsy or electrocautery-assisted resection, and formalin-fixed and paraffin-embedded (FFPE) for routine histopathology with residual tissue from the same FFPE blocks subsequently used for study-specific imaging and additional histopathological analyses.

### 1.3 Equipment

- CERO 3D Incubator and Bioreactor (OMNI Life Science, https://www.ols-bio.com/3d-cell-culture-bioreactor-cero)
- CERO tubes (OMNI Life Science, #2800005)
- QuantStudio 7 Flex Real Time PCR System (Thermo Fisher, #25008528)
- Thunder DMi8 Widefield (Leica, #25013617)
- Confocal Stellaris 5 (Leica, #25023819)
- Nikon CrestOptics Spinning Disk Confocal Microscope (Nikon) with 20X air, 0.8 NA, CFI Plan Apochromat Lambda D objective (Nikon)
- FLUOVIEW FV4000 Confocal Laser Scanning Microscope (Olympus) with 40X oil objective, X-Apo,1.4 NA (Olympus, #AMEP4908)
- 3D Sunflower Mini-Shaker (biosan, #BS-010151-AAG)
- Leica UC7 (Leica Mikrosystems, serial number: 614293)
- Formvar Support Film Slots, 2×1 mm Cu grids (Electron Microscopy Sciences, #FF2010-CU-50)
- JEOL JEM-2100Plus transmission electron microscope (JEOL Ltd., serial number: EM1753001310131)

### 1.4 Softwares

- FIJI / ImageJ 1.54p (https://imagej.net/ij) (Schindelin et al., 2012)
- Imaris v10.1 (Oxford Instruments, https://imaris.oxinst.com)
- QuantStudio Real-Time PCR System (Thermo Fisher, https://www.thermofisher.com/ch/en/home/global/forms/life-science/quantstudio-6-7-flex-software.html?erpType=Global_E1)
- Omero microscopy data repository (https://www.openmicroscopy.org)
- Anaconda Distribution (https://www.anaconda.com)
- Python 3.14.6 with Jupyter Notebook(1.1.1), Napari 0.8.0 (https://napari.org) and µ-SAM 1.0 plugins (Archit et al., 2025)
- SerialEM (David Mastronarde, University of Colorado Boulder, https://bio3d.colorado.edu/SerialEM/) (Mastronarde, 2005)
- IMOD (David Mastronarde, University of Colorado Boulder, https://bio3d.colorado.edu/imod/)
- MAGE3D (Jean Daraspe, University of Lausanne, https://wp.unil.ch/emf/software-mage3d/)

### 1.5 Scripts

- Organoid segmentation script (https://github.com/vishwa-vasani/apical-out-hBO-brightfeld-image-analysis/blob/main/organoid-segmentation.ipynb)
- Organoid size adaptive thresholding and plotting script (https://github.com/vishwa-vasani/apical-out-hBO-brightfeld-image-analysis/blob/main/adaptive-thresholding-and-plotting.ipynb)

### 1.6 Ethical statement

Participants were recruited at the Department of Neuro-Urology, Balgrist University Hospital, University of Zurich, Switzerland. The study was approved by the Cantonal Ethics Committee Zurich (BASEC Nr. 2010-0207 and 2023-00743), and all participants provided written informed consent.

### 1.7 Reagents set-up

#### 1.7.1 Fetal bovine serum (FBS) heat inactivation

FBS is stored at −20°C and thawed overnight at 4°C prior to heat inactivation. The thawed FBS is put at 56°C for 30 minutes with regular swirling (every 5 minutes). After heat-inactivation, the FBS is immediately put on ice before aliquoting. Aliquots are stored at −20°C. Thawed aliquots can be kept at 4°C for up to four weeks.

#### 1.7.2 H-6621 medium

Complete epithelial cell medium is prepared according to manufacturers’ protocol. Although the supplier recommends 5% of FBS for the H-6215 cells (as supplied with the growth factor kit), a total of 10% performed better in our hands. We generally omitted the supplied antibiotic-antimycotic solution during cell expansion – this is especially important if regular mycoplasma testing with antibiotic-antimycotic sensitive techniques is performed. The full medium is stored at 4°C for up to two months and will be hereafter referred to as H-6621 medium.

#### 1.7.3 HBO medium

HBO medium is prepared by complementing DMEM/F-12, GlutaMAX™ supplement with all the reagents outlined in Table 1 (Tomasek et al., 2026). Full medium is stored at 4°C protected from light for up to two months.

**Table 1:** HBO media preparation.

| **Reagent** | **Final Concentration** |
|---|---|
| Human FGF-10 Recombinant Protein, PeproTech® | 100 ng/mL |
| Human FGF-7 (KGF), Recombinant Protein, PeproTech® | 100 ng/mL |
| A 83-01 | 500 nM |
| B-27™ Supplement (50X), serum free | 2% |
| Y-27632 dihydrochloride | 10 µM |
| N-Acetyl-L-cysteine | 1 mM |
| Nicotinamide | 10 mM |
| PeproGMP® Human VEGF-165 Recombinant Protein, PeproTech® | 50 ng/ml |

#### 1.7.4 3dGRO® L-WRN Conditioned Media Supplement media supplement (L-WRN) medium

The L-WRN conditioned medium is stored at −20°C. Media is thawed overnight at 4°C in the dark before usage. Unused media should be aliquoted and stored at −20°C. After thawing refrozen aliquots, do not re-freeze again! Thawed medium can be kept for two weeks at 4°C.

##### Alternatives

L-WRN can be produced from L-WRN producing fibroblast-like cells (CRL-3276™, ATCC). If L-WRN conditioned medium is produced in-house, consider measuring Wnt3a levels with an ELISA suitable for cell culture supernatants.

#### 1.7.5 Live cell imaging solution (pH 7.4)

Live cell imaging solution is prepared by diluting the compounds outlined in Table 2 in ddH_2_O (Tomasek et al., 2026).

**Table 2:**
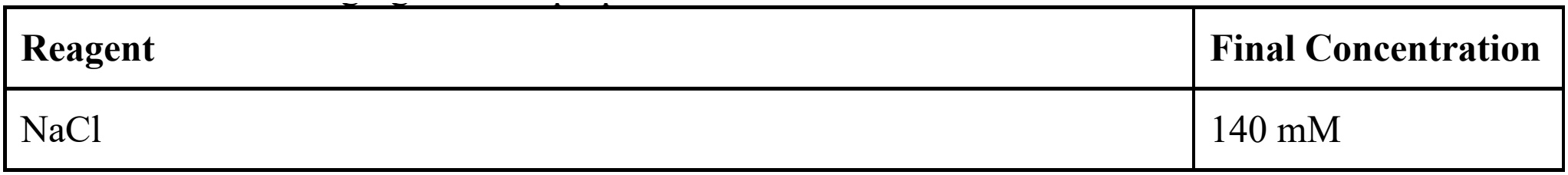
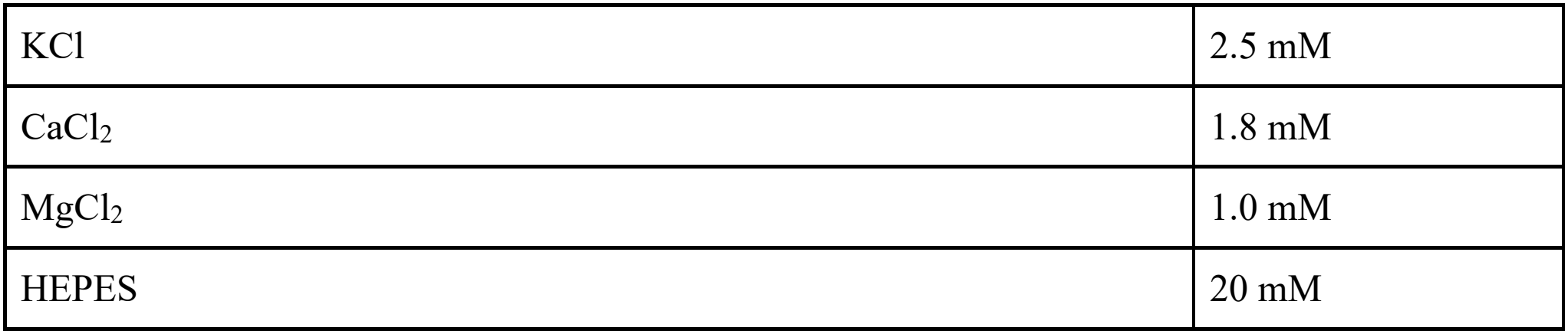
Live cell imaging solution preparation.

#### 1.7.6 Cultrex Reduced Growth Factor Basement Membrane Extract, Type 2, Pathclear (Matrigel)

The Matrigel stock is stored at −80°C upon delivery and thawed overnight at 4°C. Aliquot thawed Matrigel in pre-chilled Eppendorf tubes and store aliquots at −20°C. Defrost aliquots at 4°C for at least two hours before usage. After thawing frozen aliquots, do not freeze again! Thawed aliquots can be stored at 4°C for up to one week. Keep thawed Matrigel on ice at all times to prevent premature polymerization.

#### 1.7.7 Freezing medium

Freezing media is prepared by mixing 90% of heat-inactivated FBS with 10% of DMSO. Freezing media can be stored at 4°C for up to one week.

#### 1.7.8 4% Paraformaldehyde (PFA)

4% PFA is prepared by diluting a16% solution in phosphate-buffered saline (PBS). The solution is always prepared fresh.

#### 1.7.9 2.5% glutaraldehyde

2.5% glutaraldehyde solution is prepared from a 50% stock solution by diluting in PB buffer. The solution is always prepared fresh.

#### 1.7.10 1% bovine serum albumin (BSA) in PBS

1% w/v of BSA is added to PBS. The solution is then stirred gently until the BSA is fully dissolved. Upon filtration using a 0.22 µm filter, it can be store at 4°C for up to six months.

#### 1.7.11 Pooled human urine (PHU)

Gender PHU from BioIVT is thawed upon arrival and aliquoted. Aliquots are stored at −20°C. Thawed aliquots can be stored at 4°C for up to one week. Any precipitate forming should be brought back into solution by warming up to 37°C and vortexing rigorously.

##### Alternatives

With the necessary ethical approval, PHU can also be sampled from healthy volunteers. PHU should be filtered using a 0.22 µm filter before aliquoting and freezing.

##### Note

Urine osmolarity varies substantially over the course of the day (Brooks and Keevil, 1997) and can influence epithelial integrity and, consequently, infection dynamics (Paduthol et al., 2026). Mid-day urine typically reflects intermediate osmolarity, whereas morning urine is generally more concentrated and fluid intake can result in diluted, low-osmolarity urine. Researchers should select or standardize urine osmolarity according to the experimental question. For this study, we used urine with intermediate osmolarity (~400 mOsm/kg) as provided by BioIVT.

#### 1.7.12 Reynolds lead citrate

Reynolds lead citrate stain is prepared by mixing solution A (1.33 g lead nitrate in 15 ml H_2_O) and solution B (1.76 g sodium citrate dihydrate in 15 ml H_2_O) for 30 min in the dark (a milky precipitate will appear). 8 ml of NaOH 1N and 12 ml H_2_O are then added to solubilize the precipitate. The solution is filtered using a 0.22 µm filter, aliquoted in syringe (without air) and stored in the dark at 4°C.

#### 1.7.13 Epon resin

Mix 194 g of Embed 812 with 222 g of NMA (mix A), and separately, 156 g of Embed 812 with 136 g of DDSA (mix B). Add mix A and mix B together and homogenize.

High-BDMA Epon (used only for the very last incubation) is prepared by adding 0.26 g of BDMA in 10 g of the final mix.

## Methods

### 1.8 Step-by-step method for the production of apical-out human bladder organoids from primary cells

#### 1.8.1 H-6215 cell expansion

Duration: up to 2 weeks (depending on the initial passage number)

##### Note

H-6215 cells were routinely treated with Mycoplasma Elimination Reagent for the first expansions.

1. Start a culture of H-6215 human bladder epithelial cells in H-6621 medium according to the manufacturer’s recommendations and immediately treat with Mycoplasma Elimination Reagent for the first 3 passages
  a. *Note:* H-6215 cells can be treated for 1 passage with reagent 1 and for 2 passages with reagent 2 before omitting treatment
2. Expand the cells according to the manufacturer’s recommendations until maximum passage number of 6-8
  a. *Note:* routinely test for Mycoplasma contamination using the appropriate test kit

##### Note

Make sure to have one million cells before the next step.

#### 1.8.2 Generation of apical-out human bladder organoids in the CERO 3D Bioreactor

Duration: 8 days

##### Note

The protocol below describes the preparation of a single CERO tube for the growth of human bladder organoids from primary H-6215 cells (Figure 1A).

**Figure 1:**
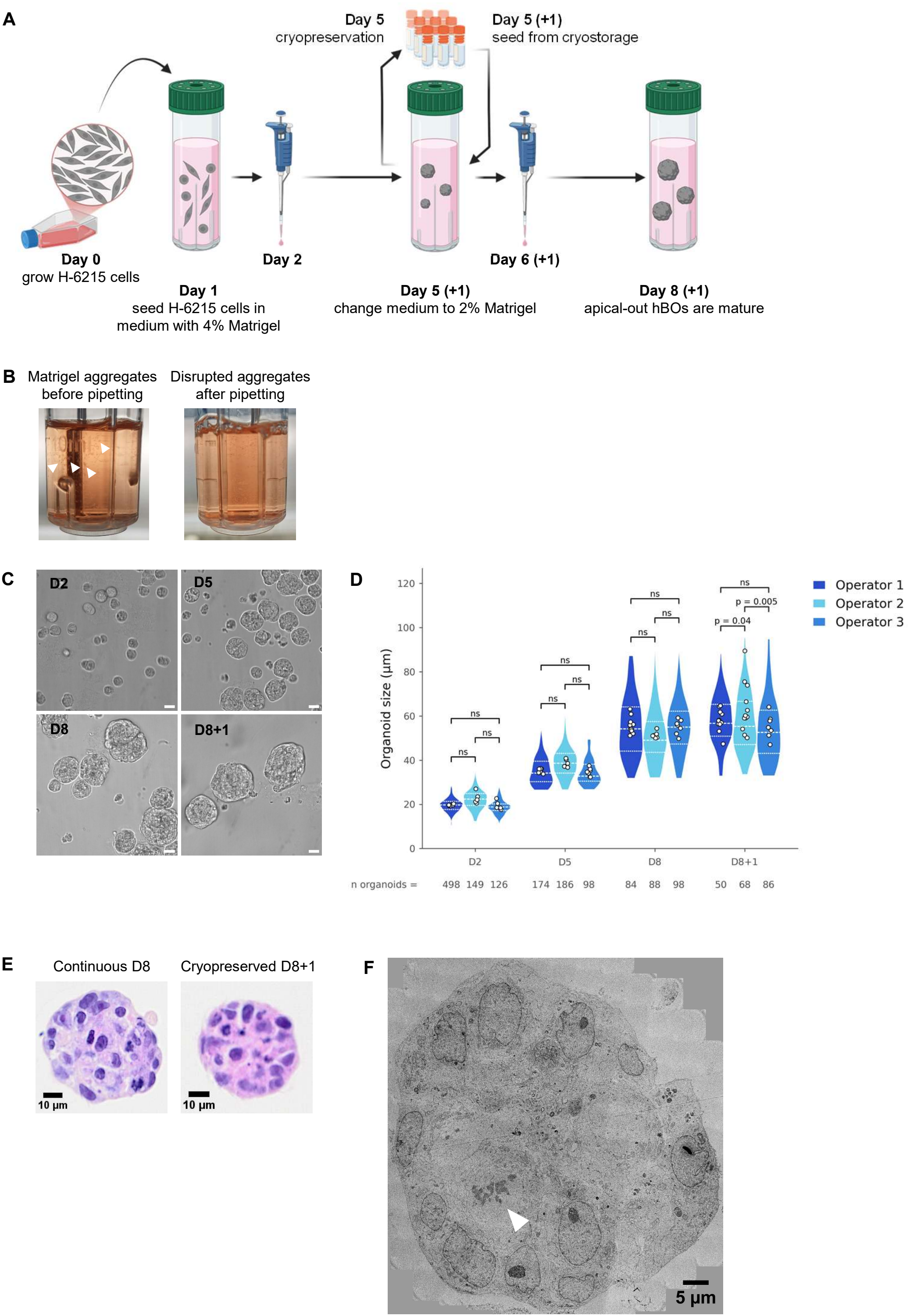
Development of apical-out human bladder organoids in suspension culture. A. Schematic of apical-out human bladder organoid generation. On day 1, one million H-6215 cells in warm HBO medium are seeded into CERO tubes with cold L-WRN containing final 4% of Matrigel. On day 2, Matrigel aggregates are broken down by pipetting. On day 5, medium is changed to fresh warm HBO medium and cold L-WRN containing final 2% of Matrigel. Here, organoids can be cryopreserved or grown further. On day 6 (or day 6+1 for cryopreserved organoids), Matrigel aggregates are again broken down by pipetting. On day 8(+1) organoids are mature and can be used for downstream applications. Created in BioRender. McKinney, J. (2027) https://BioRender.com/hw1y00y. B. On day 2 and day 5(+1), Matrigel aggregates are disrupted by pipetting. Before pipetting, aggregates are clearly visible (left; some aggregates indicated with arrow heads). Following accurate pipetting, the culture should appear uniformly turbid indicating adequate disruption of the aggregates (right). C. Representative brightfield images of organoids on days 2 (D2), 5 (D5) and 8 (D8) in continuous culture, and of organoids frozen on day 5, thawed and cultured for 3 additional days (D8+1). Scale bar 20 µm. D. Organoid size quantification using brightfield images. Each condition was determined to be either unimodal or bimodal (SI Figure 1D). The threshold was then set at the valley between the two fitted components (D5, 26.6 µm; D8, 31.7 µm; D8+1, 32.5 µm), and only objects above it were classified as organoids and analyzed. Objects smaller than 10 µm in hydraulic diameter were excluded as debris. D2 was unimodal, so all objects were included. Mean organoid diameter per image (n = number of images per group) was compared by two-way ANOVA (factors: condition and operator, with interaction), followed by Tukey’s multiple comparisons test between operators within each condition. Data shown as mean ± SD; ns, not significant. E. Representative H&E (Hematoxylin & Eosin) images of continuous (D8) and cryopreserved (D8+1) organoids. Organoids display a solid, spheroid-like shape with stratified epithelial architecture. F. TEM (transmission electron microscopy) image of a continuously grown (D8+1) apical-out hBO (6,000x magnification). The arrowhead points to a mitotic event in a dividing cell.

**Day 1: Starting of a CERO tube culture** (duration: 3 hours, hands-on time: 1 hour 40 minutes)

1. Media preparation (duration: 2 hours, hands-on time: 30 minutes)
  a. Thaw Matrigel aliquot at 4°C for at least 2 hours
  b. Thaw L-WRN aliquot at 4°C for at least 2 hours
  c. Warm up DMEM/F12 GlutaMAX™ supplement and add 10% FBS
  d. Warm up TrypLE Express
  e. Warm up 7 ml of HBO media and add Primocin at 200 µg/ml
  f. To 6 ml of cold L-WRN, add 8% Matrigel and keep on ice
2. CERO 3D Bioreactor set-up (duration: 1 hour, hands-on time: 10 minutes)
  a. Make sure the CERO 3D is connected to a CO_2_ supply and turn it on according to the manufacturer’s instructions
  b. Set heating to 37°C and the CO_2_ input to 5%.
  c. The machine should be on for at least one hour before proceeding to step 3
  d. Program a new experiment in the CERO 3D according to the manufacturer’s instructions. Set up the following parameters in the “culture” phase:
    - Rotation speed: 80 rpm
    - Rotation period: 1 sec
    - Agitation pause: none
    - Rotation pause: none
    - Agitation period: 0
    - Protocol duration: infinite
    - Omit “seeding” and “harvest” phase of the program during the run
3. Cell seeding (duration: 1 hour, hands-on time: 1 hour)
  a. Detach H-6215 cells from culture flask using TrypLE Express
    - Remove cell culture medium from the cell-culture flask
    - Rinse cells once with excess of PBS
    - TrypLE cells for maximum 10 minutes at 37°C with frequent tapping of the cell culture flask to facilitate de-attachment
    - Collect cells in a 15 ml falcon and deactivate TrypLE with excess amount of warm DMEM/F12 GlutaMAX™ supplement with FBS
    - Spin cells for 5 minutes at 200 g
    - Carefully remove the supernatant
    - Wash cells once more by adding excess amount of warm DMEM/F12 GlutaMAX™ supplement with FBS
    - Spin cells for 5 minutes at 200 g
    - Carefully remove supernatant
    - Resuspend cells in 1 ml of warm HBO media
    - Count viable cells following standard protocols using Trypan Blue staining
  b. Resuspend 1*10^6^ cells in 6 ml of the warm HBO media
  c. Put 6 ml of cold L-WRN/Matrigel in the CERO tube
  d. Add 6 ml of warm HBO/cell mix
  e. Place the CERO tube in the CERO 3D bioreactor
    - Select the respective slot and the experiment prepared earlier (step 2)
    - *Note*: before starting the experiment, unselect the “seeding” and “harvest” phase
  f. Start the program and close the CERO 3D bioreactor

##### Critical

It is essential to combine a warm cell-containing HBO medium with a cold Matrigel-containing medium during setup to enable controlled micro-aggregation of Matrigel into small droplets. This temperature contrast promotes spontaneous formation of Matrigel droplets, to which epithelial cells attach and subsequently self-organize into stratified organoids (Parigoris et al., 2021). In practice, cells are resuspended in pre-warmed medium, while Matrigel is prepared in ice-cold L-WRN conditioned medium before mixing.

##### Note

The final Matrigel concentration is 4% and the final Primocin concentration is 100 µg/ml.

**Day 2: Breaking the Matrigel clump** (duration: 5 minutes, hands-on time: 5 minutes)

##### Critical

For this and all following steps it is critical to coat all laboratory plastics (e.g. pipette tips, serological pipettes, Eppendorf tubes, falcons, …) which will get in contact with the organoids with 1% BSA solution. This step is critical to minimize organoid loss due their adhesion to plastics!

1. Break Matrigel clumps surrounding organoids
  a. Pipette organoids up and down 20 times using a 1000 µl tip
    - *Note*: restrict the pipetting volume to ~300-500 µl while pipetting to improve mechanical shear and cause a more effective dispersion of the Matrigel clumps
2. Place tube back into the CERO 3D bioreactor

##### Note

This step is required to achieve a better single-organoid distribution within the medium. Following this step, the culture should appear uniformly turbid, indicating adequate disruption of the previously clearly visible matrix aggregates (Figure 1B).

**Day 5: Media change** (duration: 3 hours, hand-on time: 45 minutes)

##### Critical

It is critical to coat all laboratory plastics with 1% BSA solution to minimize organoid loss.

1. Media preparation (timing: 2 h, hands-on: 30 minutes)
  a. Thaw Matrigel aliquot at 4°C for at least 2 hours
  b. Thaw L-WRN aliquot
  c. Warm up 6 ml of HBO media and add Primocin at 200 µg/ml
  d. To 6 ml of cold L-WRN, add 4% Matrigel and keep on ice
2. Collect organoids from CERO tube
  a. Transfer organoids from CERO tube to a 15 ml falcon tube
  b. Spin organoids at 200 g for 5 minutes
  c. Carefully remove the supernatant
  d. Gently resuspend the organoids in 6 ml of warm HBO media
3. Put 6 ml of cold L-WRN/Matrigel mix into the same CERO tube that was used before
4. Add the 6 ml of warm HBO/organoid mix
5. Place CERO tube back in the CERO 3D bioreactor

##### Note 1

From day 5 onward, the final Matrigel concentration is reduced to 2% while Primocin stays at 100 µg/ml. If an infection assay is planned with the matured organoids, Primocin has to be omitted from this step onward.

##### 1.8.2.1 Optional

Cryopreservation (duration: 25 hours, hands-on time: 30 minutes)

###### Critical

It is critical to coat all laboratory plastics with 1% BSA solution.

Instead of changing media on day 5, it is possible to cryopreserve the organoids for later usage. Depending on the downstream application, all organoids or a portion can be cryopreserved (e.g. we have successfully frozen 95% of the organoid culture on day 5 and kept only 5% growing, resulting in roughly 10,000-15,000 organoids on day 8 for experimental purpose).

1. Collect organoids from CERO tube
  a. Transfer organoids from CERO tube to a 15 ml falcon tube
  b. Spin organoids at 200 g for 5 minutes
  c. Carefully remove the supernatant
2. Freeze organoids
  a. Gently, but quickly resuspend organoids in up to 12 ml of freezing media
  b. Aliquot 500-1000 µl per cryovial
  c. Quickly freeze aliquots at −80°C using a Mr. Frosty
  d. 24 hours later, transfer cryovials to liquid nitrogen for prolonged storage

##### 1.8.2.2 Optional

Thawing (duration: 4 hours, hands-on time: 45 minutes)

It is alternatively possible to start a culture from day 5-cryopreserved organoids.

###### Note

When thawing an aliquot of the cryopreserved organoids, the total media volume per CERO tube can be decreased to 6 ml.

###### Critical

It is critical to coat all laboratory plastics with 1% BSA solution.

1. Media preparation (timing: 2 h, hands-on: 30 minutes)
  a. Thaw Matrigel aliquot at 4°C for at least 2 hours
  b. Thaw L-WRN aliquot 4°C for at least 2 hours
  c. Warm up DMEM/F12 GlutaMAX™ supplement
  d. Warm up 3 ml of HBO media and add Primocin at 200 µg/ml
  e. To 3 ml of cold L-WRN, add 4% Matrigel and keep on ice
2. Thaw organoids
  a. Quickly thaw cryovial in a warm bead bath
  b. Transfer thawed organoids to a 15 ml falcon tube using a 1000 µl tip
  c. Slowly but continuously add excess volume of warm DMEM/F-12, GlutaMAX™ supplement media
  d. Spin at 200 g for 5 minutes
  e. Carefully remove the supernatant
  f. Gently resuspend organoids in 3 ml of warm HBO media
3. Put 3 ml of cold L-WRN/Matrigel mix into the CERO tube
4. Add the 3 ml of warm HBO/organoid mix
5. Place CERO tube in the CERO 3D bioreactor

###### Note

The final Matrigel concentration is 2% while Primocin stays at 100 µg/ml. If an infection assay is planned with the matured organoids, Primocin has to be omitted from this step onward.

**Day 6 (or thaw day + 1): Breaking the Matrigel clump** (duration: 5 minutes, hands-on time: 5 minutes)

###### Critical

It is critical to coat all laboratory plastics with 1% BSA solution.

1. Break Matrigel clumps surrounding organoids
  a. Pipette organoids up and down 20 times using a 1000 µl tip
    - *Note*: restricting the pipetting volume to ~300-500 µl while pipetting to improve mechanical shear and cause a more effective dispersion of the Matrigel clumps
2. Place tube back into the CERO 3D bioreactor

###### Note

This step is required to achieve a better single-organoid distribution within the medium. Again, following this step, the culture should appear uniformly turbid, indicating adequate disruption of the previously clearly visible matrix aggregates (Figure 1B).

**Day 8**: **Apical-out hBOs are ready for downstream applications** (duration: variable)

###### Note

To avoid shear stress on organoids, cut small pipette tips (≤ 200 µl) with sterile scissors for small volume applications.

### 1.9 Fixation of organoids

#### Critical

It is critical to coat all laboratory plastics with 1% BSA solution to avoid organoid loss.

1. Fixation of organoids
  a. Transfer organoids from CERO tube to a 15 ml falcon tube
  b. Spin at 200 g for 5 minutes
  c. Carefully remove the supernatant
  d. Fix organoids in a small volume of 4% PFA at 4°C overnight
  e. Wash by adding excess amount of PBS
  f. Spin at 200 g for 5 minutes
  g. Carefully remove the supernatant
  h. Store organoids in PBS at 4°C or proceed

### 1.10 Measuring size distributions of organoids over development

#### Note

Organoid size was measured by using extended depth of field (EDF) script in the SWIFT (Single-organoid Workflow for quantitative Imaging classiFication and Tracking) pipeline (Bracq et al., 2026) and using a custom code for segmentation and statistical analysis (scripts provided with this publication). This experiment is performed with fixed organoids (see section 1.9).

#### Critical

It is critical to coat all laboratory plastics with 1% BSA solution to avoid organoid loss prior to imaging.

1. Sample preparation
  a. Pipette fixed organoids up and down 10 times to break up any clumps with 1 ml tip
  b. Transfer 100 µl of fixed organoids from refrigerated storage into 96 well µ-plate (ibidi)
  c. Repeat 3 more times to prepare 4 wells per sample
  d. Add 100 µl of 1% BSA solution to prevent organoid clumping by reducing nonspecific adhesion *Note*: Allow organoids to settle to the bottom over 30 minutes before imaging.
2. Widefield imaging
  a. On the Nikon CrestOptics microscope, utilize the spinning disk confocal mode in brightfield imaging setting
  b. Locate a suitable field of view that captures several organoids within a single frame, *Note:* image a field of view where the least number of organoids are vertically on top of another for accurate segmentation
  c. Record a z-stack with a z-spacing of 2 µm over a z-range in which all organoids within the field of view are captured
  d. Load all image files onto an Omero repository
3. Image analysis
  a. Load and run the EDF script in FIJI
  b. Within the pop-up dialogue box, specify the Omero login credentials as well as the dataset ID for the repository containing brightfield images
  c. Allow the script to complete executing. This script will look through each z plane within a z-stack, and project each in-focus organoid from its respective z plane onto a single 2D plane
  d. Utilize the projected 2D images for subsequent automatic segmentation and morphological measurement
  e. From the Anaconda power shell, navigate to the environment where Napari and µ-SAM are installed. Load the segmentation and size analysis Jupyter notebook
  f. Run the Jupyter notebook to see raw segmented images. Cycle through all images and manually remove any incorrect organoid segments by following instructions on the screen
  g. After all the organoid segments have been verified, allow the Jupyter notebook to finish executing to generate the measurement summary Excel sheet
  h. To separate organoids from small cell clusters, determine a size threshold independently for each condition by fitting one- and two-component Gaussian mixture models to log-transformed diameters. A condition can be considered bimodal when the two-component model fit better (ΔBIC > 10) and the two peaks are well separated (Ashman’s D > 2)
  i. In case of bimodal distribution, define size threshold as the diameter at which the two Gaussian curves intersect, i.e. an organoid has equal probability of belonging to either curve. Only consider organoids above this size threshold for further analysis.

### 1.11 RNA extraction and qPCR for gene expression evaluation

#### Note

For this protocol, RNA of one day old organoids grown from H-6215 cells in the CERO 3D (D1) served as control.

#### Critical

It is critical to coat all laboratory plastics with 1% BSA solution to avoid organoid loss prior to RNA extraction.

1. RNA extraction
  a. Transfer organoids from CERO tube to a 15 ml falcon tube
  b. Spin at 200 g for 5 minutes
  c. Carefully remove the supernatant
  d. Follow standard RNA extraction procedure and keep good practice handling RNA
    - *Note*: We used the RNAse Easy Mini Kit following manufacturer’s protocol. As recommended, the lysis buffer was supplemented with 1% β-mercaptoethanol to prevent RNA degradation.
2. cDNA conversion Convert equal amounts of RNA for each sample to first-strand cDNA using a reverse transcriptase (e.g. SuperScript IV Reverse transcriptase). Follow manufacturer’s protocol.
  - *Note*: We used 935 ng of total RNA based on the lowest RNA concentration measured in one of our samples.
3. RT-qPCR reaction Use a universal master max with Taq polymerase (e.g. SYBR™ Green Universal Master Mix). Follow manufacturer’s protocol in setting-up a standard RT-qPCR reaction for each target gene in technical duplicates. Use 1 µl cDNA per reaction and 500 nM primers (Table 3).

**Table 3.**
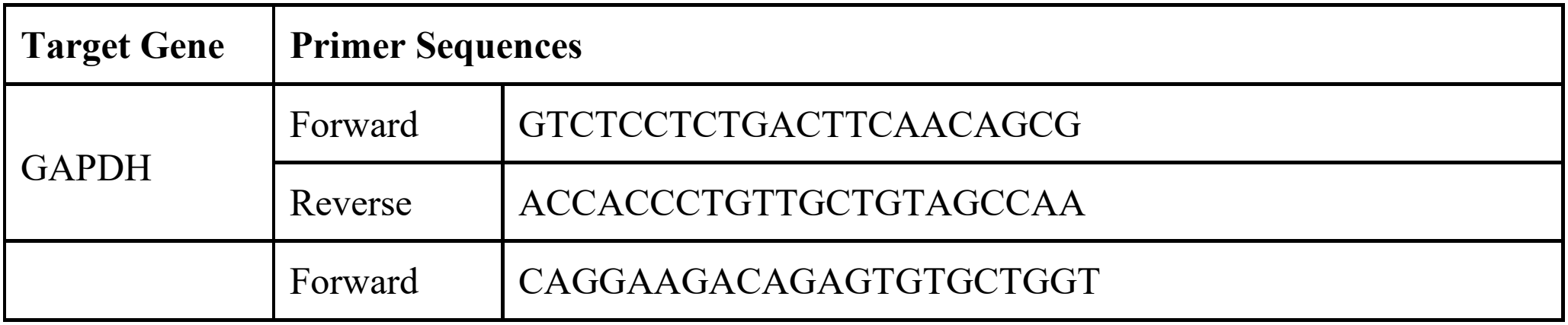
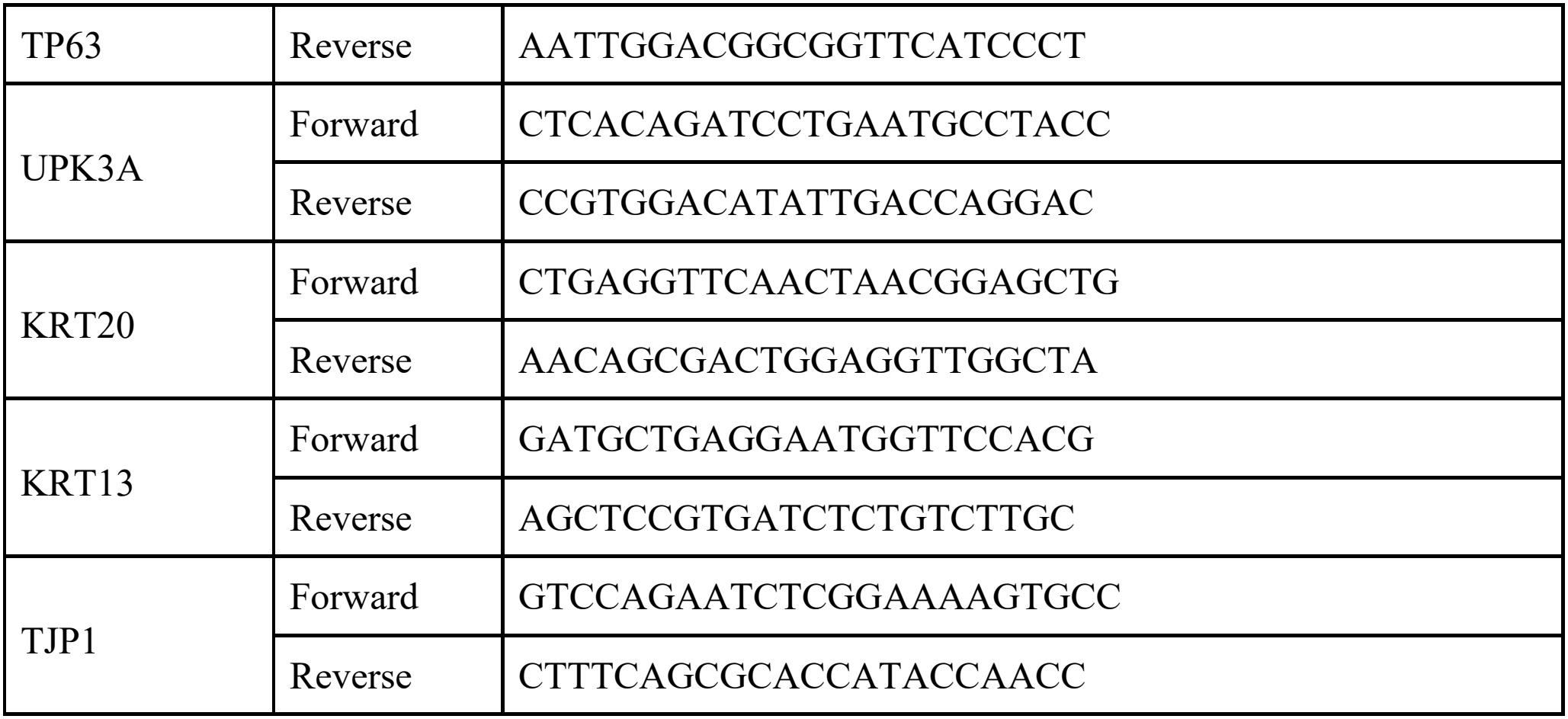
qPCR primers.

### 1.12 TEM imaging of organoids for characterization

#### 1.12.1 Preparation of TEM sections

1. Fixation and post-fixation
  a. Fix organoids in glutaraldehyde (2.5% in PB buffer) for 1 hour at RT, then wash twice in PB buffer
  b. Spin down organoids (3000 rpm) and remove supernatant, then post-fix organoids in osmium tetroxide 1% with 1.5% of potassium ferrocyanide in PB buffer
2. Embedding and resin infiltration
  a. Spin down organoids (3000 rpm) and remove supernatant, then embed organoids in 2% agarose in water
  b. Dehydrate the organoids in acetone solution at graded concentrations: 30% acetone for 40 minutes, 70% acetone for 40 minutes, 100% acetone 2x for 1 hour
  c. Infiltrate in Epon resin at graded concentrations: Epon 33% in acetone for 2 hours, Epon 66% in acetone for 4 hours, Epon 100% in acetone 2x for 8 hours). The last incubation should be done in Epon of higher BDMA concentration
  d. Polymerize for 48 hours at 60°C in an oven
3. Sectioning and grid preparation
  a. Cut 50-nm thick sections using a Leica UC7 ultramicrotome
  b. Pick up sections on a Formvar-coated grid 2×1 mm
4. Post-staining
  a. Post-stain sections with uranyl acetate (4% in H_2_O) for 10 minutes, rinse several times with H_2_O, then incubate in Reynolds lead citrate in H_2_O for 10 minutes, rinse several times with H_2_O at the end

#### 1.12.2 TEM imaging

1. Acquire TEM images
  a. On a JEOL JEM-2100Plus transmission electron microscope equipped with a TVIPS TemCamXF416 digital camera (TVIPS GmbH, Gauting, Germany), set at an acceleration of 120 kV, take tiled images at the desired magnification (here, 6000x) using the Serial EM software package
  b. Align tiles images with the MAGE3D program using the software IMOD

### 1.13 Staining of organoids and formalin-fixed and paraffin-embedded (FFPE) bladder tissue sections for characterization

#### 1.13.1 Processing and staining of paraffin-embedded sections

##### Note

Bladder biopsies samples were already obtained as FFPE and 4 µm cut samples on slides.

1. Embedding and slicing of organoids
  a. Warm up HistoGel to 65°C till softened
  b. Remove any residual PBS from fixed organoids by spinning at 200 g for 5 minutes
  c. Resuspend organoids in minimum of 50 µl warm HistoGel *Note*: Use a cut 200 µl pipette tip due to increased viscosity of HistoGel. Avoid bubble formation while pipetting and solidification of HistoGel.
  d. Transfer HistoGel/organoid mix to a Cryomold and let solidify as small hemispherical domes
  e. After solidification, transfer sample to HistoSette cassette and fix for 16-20 hours in 4% PFA
  f. Wash sample in HistoSette with excess amount of PBS
  g. Dehydration and clearing: submerge HistoGel dome 2 times in 70% Ethanol, 1 time in 95% Ethanol, 3 times in 100% Ethanol and finally 3 times in xylene, each for 30-60 minutes
  h. Paraffin infiltration and embedding: submerge HistoGel dome in 60°C warm paraffin wax and embed in paraffin on HistoSette
  i. Cut 4 µm thick slices on a microtome
  j. Place slices on microscopy slides and store at 4°C for later use or continue
2. Deparaffinization and rehydration of sample slices
  a. Remove paraffin: submerge slides 2 times for 5-10 minutes each in xylene
  b. Remove xylene and fixate: wash 2 times for 3-5 minutes each with 100% Ethanol, then 1 time each for 3 minutes with 95% and 70% Ethanol
  c. Hydrate cells: rinse for 2 minutes with water
  d. Proceed immediately or store in PBS up to maximum 1 week at 4°C before antigen retrieval
3. Antigen retrieval
  a. Prepare trisodium citrate buffer pH 6.0 and heat up inside retriever chamber to 60°C
  b. Submerge slides in trisodium citrate buffer for 5 minutes
  c. Heat-induced epitope retrieval (HIER): immerge slides in preheated trisodium citrate buffer and heat at 95°C for 20 minutes
  d. After cooldown of buffer to 60°C, rinse slides once with buffer and store in PBS
  e. Proceed immediately without letting slides dry off or store in PBS up to maximum 48 hours at 4°C before staining
4. Hematoxylin and Eosin (H&E) staining
  a. Submerge slides in hematoxylin solution for 5-10 minutes
  b. Rinse with water for 1-2 minutes; differentiate in acid alcohol for 1-2 seconds if desired and quickly rinse with water
  c. Bluing: dip the slides in ammonia water for 30-60 seconds, then rinse with water for 5 minutes
  d. Eosin staining: submerge the slides in Eosin Y for 1-2 minutes, then quickly rinse with water
  e. Dehydration and clearing: dip the slides 10 times in 95% Ethanol, then place in 100% Ethanol two times for 2-3 minutes each, followed by 2 times in Xylene for 5 minutes
  f. Add mounting media of choice, and seal with cover glass and nail polish before imaging on a slide scanning microscope (e.g. Olympus Slide Scanner)
5. Immunostaining:
  a. Blocking: after mark boundaries of paraffin sections with a hydrophobic pen, block marked areas in 1% BSA solution for 30-60 minutes at RT
  b. Primary antibody: stain marked areas with primary antibodies (Table 4) diluted 1:100 in 1% BSA solution overnight at 4°C *Note*: Place slides into a stain tray filled with water for overnight staining to prevent drying.
  c. Rinse excessively with PBS
  d. Secondary antibodies (Table 4): stain marked areas with respective secondary antibodies diluted 1:1000 in 1% BSA solution for 120 minutes at RT in the dark; if desired add 1:1000 DAPI
  e. Rinse three times excessively with PBS
  f. Add mounting media of choice, and seal with cover glass and nail polish before imaging on a Confocal microscope (e.g. Leica Stellaris 5) using a 63x oil immersion objective

**Table 4.**
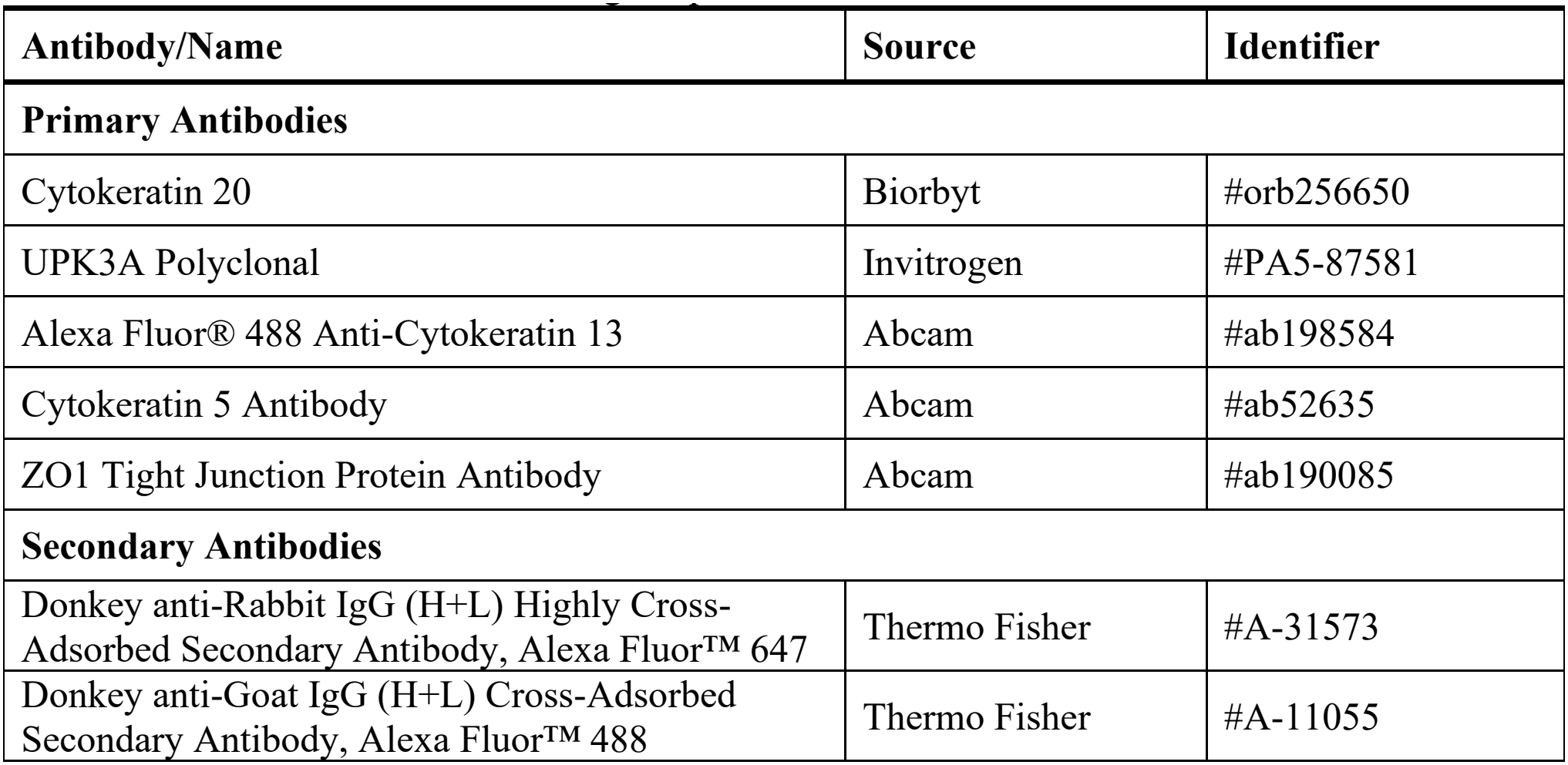
Antibodies for immunostainings of paraffin-embedded sections.

#### 1.13.2 Whole mount immunofluorescence staining

##### Critical

It is critical to coat all laboratory plastics with 1% BSA solution to avoid organoid loss.

##### Critical

When staining for membrane proteins (e.g., uroplakins), perform primary antibody incubation after blocking but prior to permeabilization. This ensures that surface-exposed proteins are detected, while intracellular or immature protein pools remain unstained. After completion of surface staining with primary antibodies, proceed with permeabilization and blocking before staining for intracellular markers.

1. Immunostaining:
  a. Take an aliquot of fixed organoids
  b. Spin at 200 g for 5 minutes and carefully remove the supernatant
  c. Permeabilize cells: add 100 µL of 0.1% triton-X in PBS for 10 minutes at RT, then wash by adding 900 µl of excess PBS followed by spinning at 200 g for 5 minutes and carefully removing the supernatant
  d. Blocking: add 200 µl 1% BSA solution for 30-60 minutes at RT
  e. Primary antibody (Table 5): directly dilute primary antibodies 1:100 into organoids blocking in BSA and stain overnight at 4°C on a rotating platform
  f. Wash with 800 µl excess of PBS, then spin at 200 g for 5 minutes and carefully remove the supernatant
  g. Secondary antibodies (Table 5): stain with 200 µl of secondary antibodies diluted 1:1000 in 1% BSA solution for 120 minutes at room temperature in the dark; if desired add 1:800 phalloidin labeling probe and 1:1000 DAPI
  h. Wash with 800 µl of excess of PBS, then spin at 200 g for 5 minutes and carefully remove the supernatant
  i. Add mounting media of choice, transfer organoids to microscopy slide and seal with cover glass and nail polish before imaging on a Confocal microscope (e.g. FLUOVIEW FV4000 Confocal Laser Scanning Microscope) using a 40x oil immersion objective

**Table 5.**
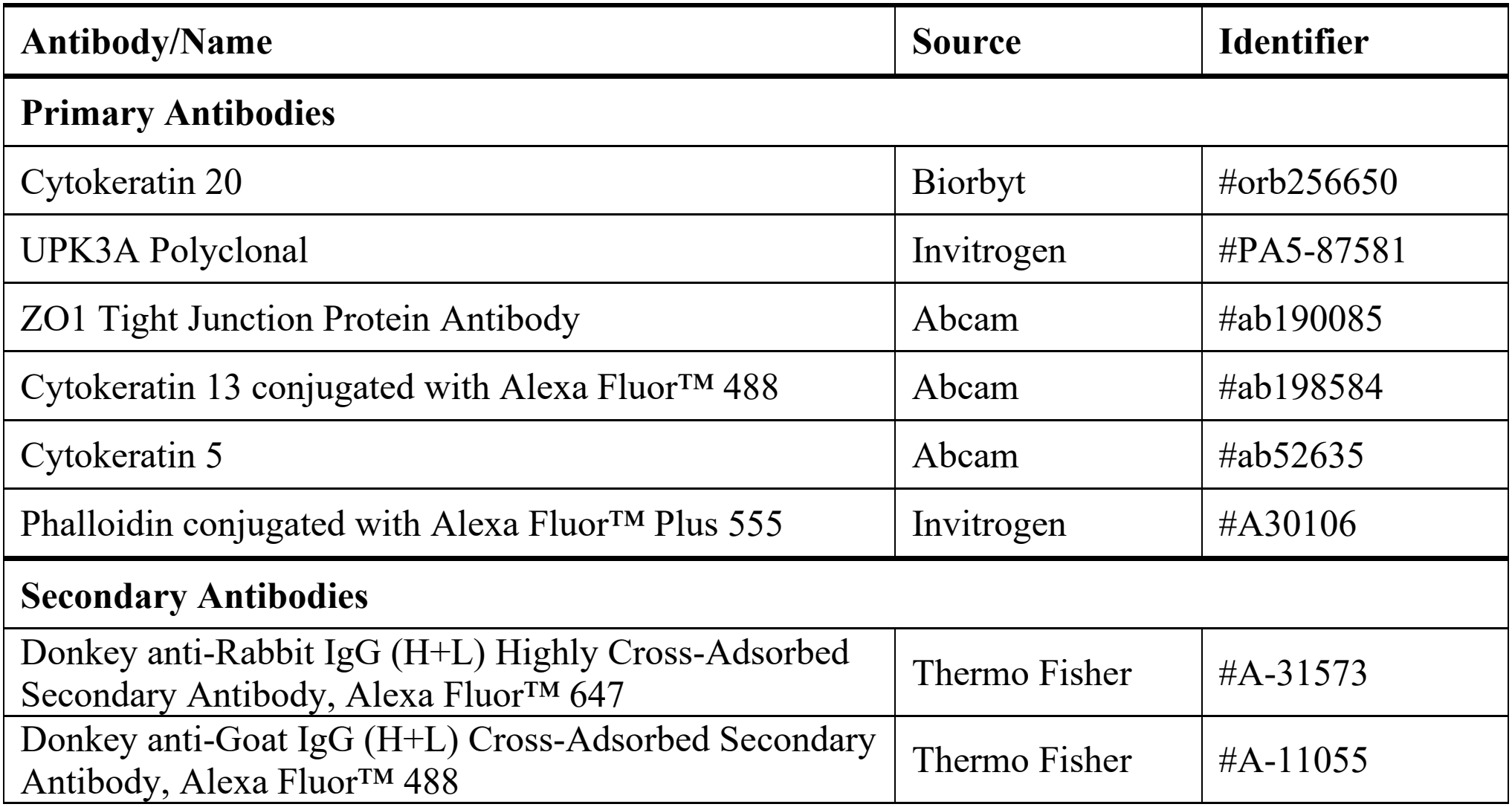
Antibodies for immunostainings of whole organoid staining.

Note: To preserve the spherical shape of organoids without compressing them under the cover glass, SecureSealTM Frame or other adhesive spacers might be used.

### 1.14 Example application

#### 1.14.1 Organoid survival in intermediate osmolarity PHU

##### Note

Cryopreserved organoids are thawed, processed and used on Day 8+1 as described in 1.6.

##### Critical

It is critical to coat all laboratory plastics with 1% BSA solution to avoid organoid loss.

1. Media preparation
  a. Warm up live cell imaging solution
  b. Thaw PHU aliquot quickly at 37°C
    - Add 1 drop of NucBlue Live ReadyProbe per mL of PHU (Hoechst 33342), 3 µM DRAQ7 dye and 1 µg/ml WGA-555
  c. Warm up 50/50 mix of HBO and L-WRN
    - To 1 ml, add 1 drop of NucBlue Live ReadyProbe (Hoechst 33342), 3 µM DRAQ7 dye and 1 µg/ml WGA-555
2. Survival time course
  d. Transfer organoids from CERO tube to two 15 ml falcon tube
  e. Spin at 200 g for 5 minutes
  f. Carefully remove the supernatant
  g. Add warm 50/50 HBO/L-WRN media or 100% PHU including Hoechst, DRAQ7 and WGA
  h. Transfer organoids to Eppendorf tubes and place on a rotating platform (e.g. 3D Sunflower Mini-Shaker) at minimum speed in a CO_2_ incubator at 37°C degree
  i. After 2 hours (time point 1) and 4 hours (time point 2):
    - Sample a 250 µl aliquot from each Eppendorf
    - Spin at 200 g for 5 minutes
    - Carefully remove the supernatant
    - Resuspend organoids in 100 µl live cell imaging solution and place in a 96-well multi-well plate (e.g. µ-plate 96 well square plate)
    - Image on a fluorescent microscope (e.g. Leica Thunder Widefield) using a 25x water immersion objective
      - Select several fields of views for each timepoint based on the WGA-555 signal of the organoids
      - Image a Z-stack with 2 µm step size to capture most of the organoids in focus
      - After deconvolution, analyze images based on their Hoechst 33342 and DRAQ7 signal per organoid
        - Perform maximum projection of images
        - Draw regions of interest around individual organoids based on WGA signal
        - Extract Hoechst 33342 and DRAQ7 signals

#### 1.14.2 Organoid infection in intermediate osmolarity pooled human urine

##### Note

The day before infection (day 7 of organoid culture), prepare an overnight culture of the UPEC strain in 3 ml 100% PHU at 37°C under static conditions. The bacterial strain should either constitutively express a fluorescent protein or be detected after fixation using an antibody against bacterial lipopolysaccharide (LPS). For the experiments presented here, we used a mutant of the UPEC laboratory strain CFT073 (Welch et al., 2002) constitutively expressing *sfGFP* from the chromosome (Simonet et al., 2022).

##### Critical

Remove all antibiotics from the organoid culture medium on the day before infection to prevent interference with bacterial growth.

##### Note

Infection can be performed in an Eppendorf tube placed in a 37°C incubator on a slow orbital shaker (e.g., 5 rpm using a 3D Sunflower Mini-Shaker) or directly in the CERO 3D bioreactor. Follow institutional biosafety regulations for all experiments involving UPEC.

1. Organoid infection:
  a. The day before infection, grow UPEC overnight in 100% PHU under static conditions
  b. The day of the infection, dilute the bacterial overnight culture 1:100 in pre-warmed 100% PHU
  c. Incubate the diluted bacterial culture for 3 hours at 37°C under static conditions
  d. Collect the organoids by centrifugation at 200 g for 5 minutes and carefully remove the supernatant
  e. Resuspend the organoids in 1 ml of bacterial solution *Note*: Under the conditions described here, the resulting bacterial concentration was approximately 1.5-1.7*10^8^ CFU/ml
  f. Transfer the organoid-bacteria suspension to an Eppendorf tube and incubate for 4 hours at 37°C with slow orbital shaking (e.g. 5 rpm orbital speed)
2. Gentamicin protection assay and post-infection incubation:
  a. Thaw Matrigel aliquot at 4°C for at least 2 hours
  b. Thaw L-WRN aliquot 4°C for at least 2 hours
  c. Prepare 50/50 HBO/L-WRN media
    - mix 0.5 ml of cold HBO with 0.5 ml of cold L-WRN
    - add Matrigel to a final concentration of 2%
    - add gentamicin to a final concentration of 10 µg/ml
    - prewarm at 37°C
  d. After 4 hours of infection, add gentamicin to the infected organoids to a final concentration of 100 µg/ml
  e. Incubate for 1 hour at 37°C with slow orbital shaking
  f. Collect the organoids by centrifugation at 200 g for 5 minutes and carefully remove supernatant
  g. Resuspend organoids in the previously prepared pre-warmed 50/50 HBO/L-WRN media supplemented with 2% Matrigel and 10 µg/ml gentamicin
  h. Incubate for an additional 16 hours at 37°C with faster shaking (e.g. 60 rpm orbital speed)
  i. Collect the organoids by centrifugation at 200 g for 5 minutes and carefully remove supernatant
  j. Fix the organoids in freshly prepared 4% PFA for 1 hour at 4°C
  k. Wash the organoids extensively with PBS and store in PBS at 4°C until further processing *Note*: The two-step gentamicin treatment is used to eliminate extracellular bacteria while maintaining gentamicin exposure during the subsequent incubation period, thereby allowing detection of bacteria that have invaded or persisted within the organoids.
3. Imaging of infected organoids:
  a. Take an aliquot of fixed organoids
  b. Spin at 200 g for 5 minutes and carefully remove the supernatant
  c. Permeabilize cells: add 0.1% triton-X in PBS for 10 minutes, then wash with excess of PBS by spinning at 200 g for 5 minutes and carefully removing the supernatant
  d. Blocking: add 1% BSA solution for 30-60 minutes at RT
  e. Stain organoids: 1:800 phalloidin labeling probe and 1:1000 DAPI in 1% BSA solution for 1 hour at RT in the dark *Note*: If the bacterial strain does not express a fluorescent protein, bacteria can be detected using an appropriate anti-LPS antibody after fixation and permeabilization.
  f. Wash with excess amount of PBS, then spin at 200 g for 5 minutes and carefully remove the supernatant
  g. Add mounting media of choice, transfer organoids to microscopy slide and seal with cover glass and nail polish before imaging on a Confocal microscope (e.g. Leica Stellaris 5) using a 63x oil immersion objective *Note*: To preserve the spherical shape of organoids without compressing them under the cover glass, SecureSealTM Frame or other adhesive spacers might be used.

## Anticipated Results

### 1.15 Morphology during organoid maturation

The protocol is reproducible across multiple independent operators of primary cells, consistently yielding stratified apical-out organoids.

Human apical-out bladder organoids form rapidly in suspension culture in the CERO 3D bioreactor, with organoid size progressively increasing over time (Figure 1C, D and SI Figure 1A-C). On Day 2, initial small multicellular aggregates that would later form larger organoids already start to form. By Day 5 there is a clear emergence of a bimodal population of several small multicellular aggregates and a smaller number of large organoids (SI Figure 1C). By day 8, this bimodal distribution is still present, and mature organoids reach an average diameter of 54.8 ± 12.3 µm, with no observable difference between continuously cultured organoids (D8) and organoids cryopreserved at day 5 and subsequently recovered for 3 additional days (D8+1).

Although a small fraction of organoids displays incorporated Matrigel aggregates of variable size within multiple cell layers (SI Figure 2A, B), the majority of organoids in both conditions yield solid, spheroid structures with a stratified epithelial architecture consisting of multiple cell layers and no visible central lumen or necrotic core (Figure 1E).

TEM imaging shows the compact architecture of apical-out hBOs and further reveals within some organoids the presence of a small fraction of presumably apoptotic cells, whereas most organoids show clearly dividing cells (Figure 1F, SI Figure 2B).

### 1.16 Validation of apical-out phenotype

RT-qPCR revealed a trend toward increased expression of urothelial differentiation markers, including UPK3A, KRT20, KRT13, and TP63, following organoid differentiation compared with day 1 organoids (Figure 2A). Although fold changes varied between independent biological replicates (different passages and operators) and did not reach statistical significance, these trends were consistent with the differentiation patterns observed by immunofluorescence and histological analysis (Figure 2B-D, Figure 3A-C).

**Figure 2:**
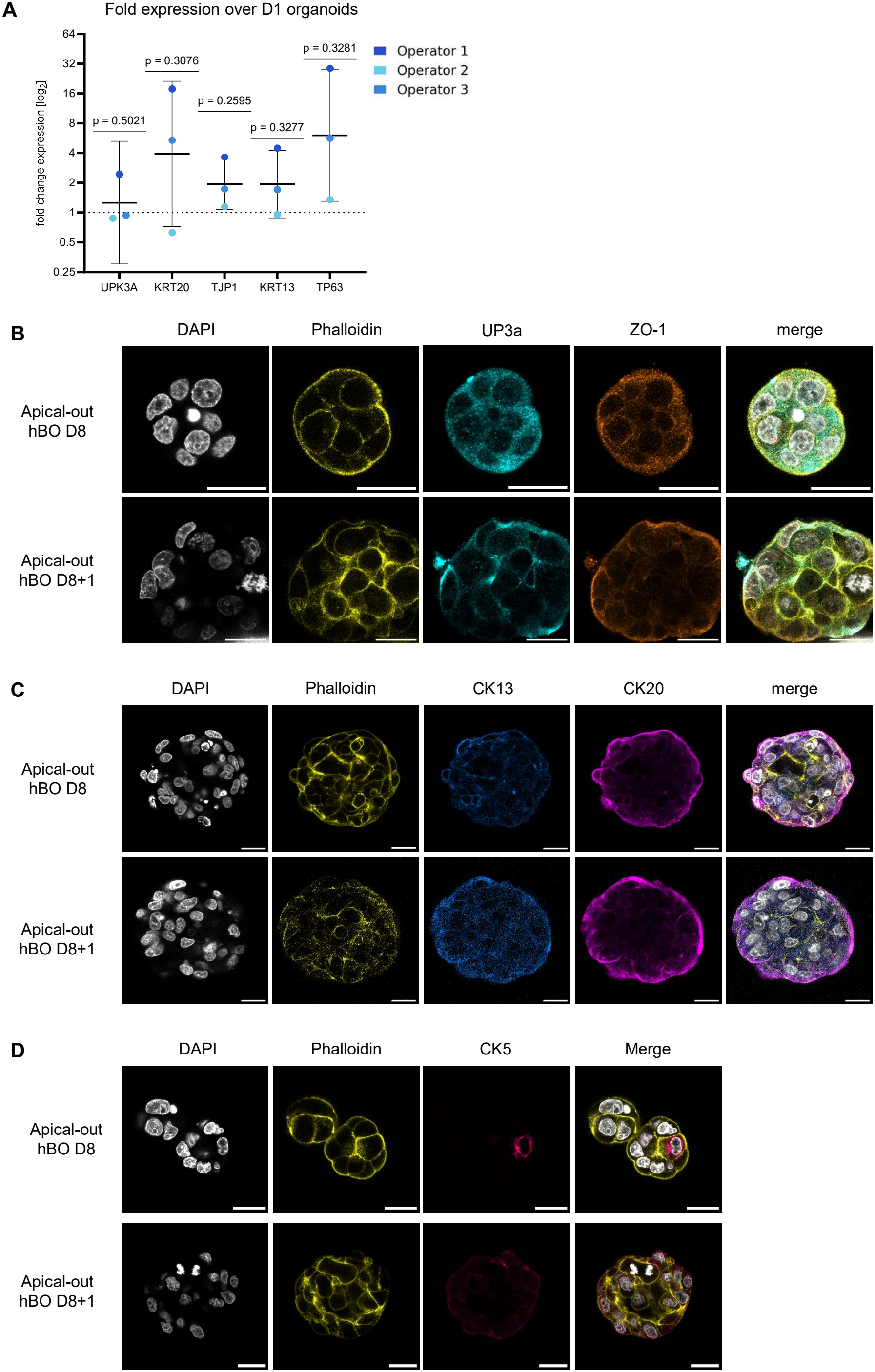
Validation of differentiation for continuous cultures (D8) and cryopreserved ones (D8+1). A. Gene expression of differentiation markers of mature apical-out organoids (D8), compared to early organoids (D1). Umbrella cell markers – cytokeratin 20 (KRT20), uroplakin 3a (UPK3A). Intermediate cell marker – cytokeratin 13 (KRT3). Progenitor cell marker – tumor protein p63 (TP63). Tight junction marker – TJP1 (ZO-1). Mean gene expression (N = 3 biological replicates) per marker was compared to the hypothetical value 1 by one sample t test. Data shown as mean ± 95% confidence interval. B-D. Whole mount immunofluorescence images (continuous and cryopreserved). Nuclei, grey. Actin, yellow. (B) CK13, blue. CK20, magenta. (C) UP3a, cyan. ZO-1, amber. (D) CK5, bright pink. Scale bar 20 µm.

**Figure 3:**
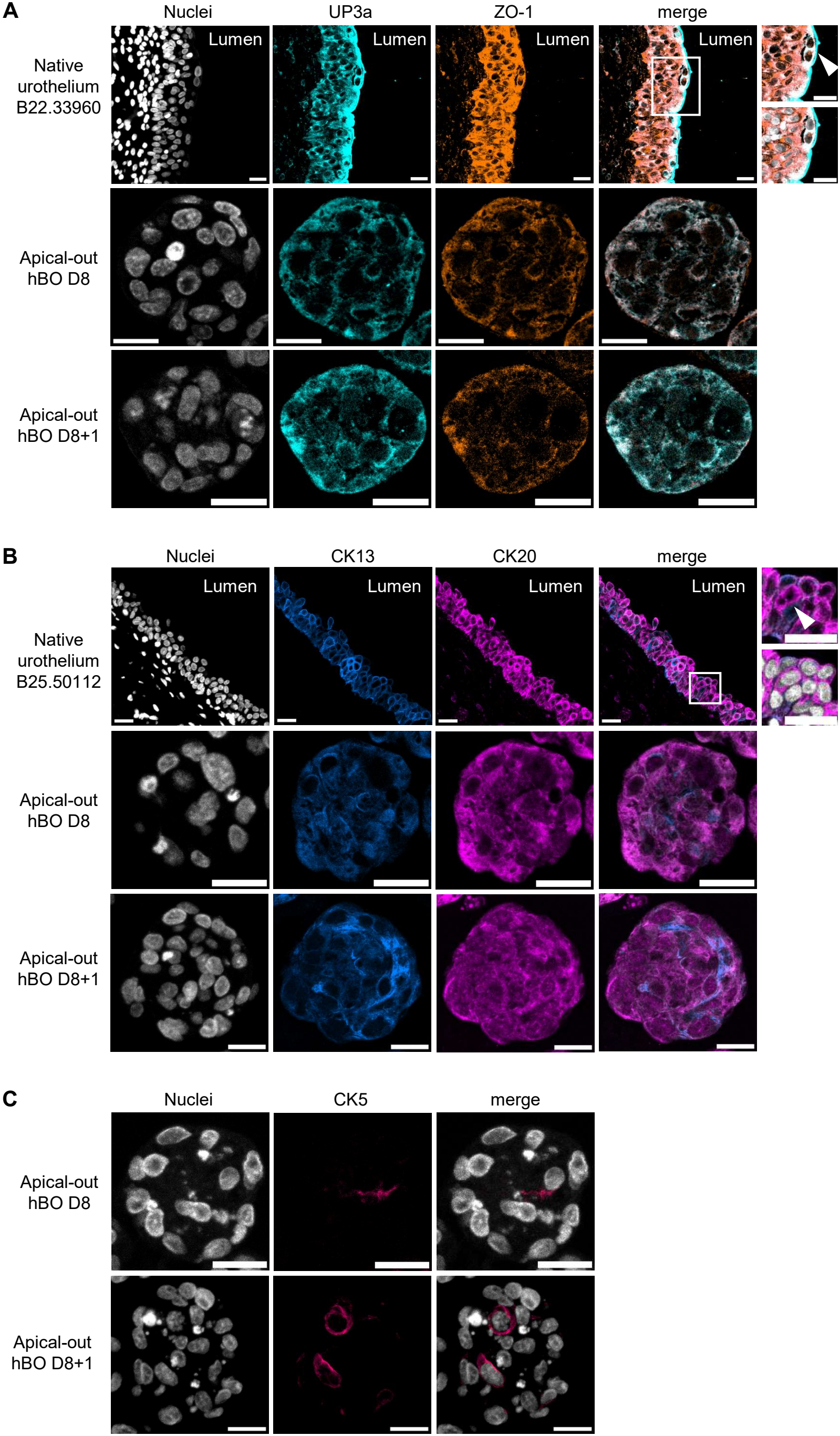
Validation of apical-out phenotype for continuous cultures (D8) and cryopreserved ones (D8+1). Immunofluorescent images of histology sections. Representative images of 4 µm sections of native urothelium and apical-out human bladder organoids. Binucleated umbrella cells indicated with arrow heads. Nuclei, grey. (A) CK13, azure. CK20, magenta. (B) UP3a, cyan. ZO-1, amber. (C) CK5, pink. Scale bar 20 µm.

Compared to day 1 organoids, TJP1 (ZO-1) transcript expression in mature organoids was increased (Figure 2A) and immunofluorescence confirmed clear junctional localization of ZO-1 in differentiated organoids (Figure 2B, 3A and SI Figure 3A, B), consistent with the formation of epithelial barrier structures on the outward-facing surface of the organoids.

Immunofluorescence and histology further confirm urothelial stratification and apical-out polarity. Whole-mount immunostaining of intact organoids and of cryosections verified spatially organized expression of differentiated umbrella cell markers at the outer surface such as of UP3a (Figure 2B, 3A and SI Figure 3A, B) and CK20 (Figure 2C, 3B and SI Figure 4A, B). Intermediate urothelial markers (CK13) were expressed within suprabasal layers (Figure 2C, 3B and SI Figure 4A, B) and basal markers (CK5) were found to be distributed through the organoid (Figure 2D, 3C and SI Figure 5A, B). This stratified organization was observed across multiple independent preparations and strongly reflects a similar pattern found in the native bladder epithelium (Figure 3A, B and SI Figure 3B, 4B). Nuclear counterstaining and membrane markers allow clear illustration of the tissue architecture and confirm the absence of a central lumen. Finally, cryopreservation (D8+1) did not alter the spatially organized expression of any observed urothelial marker.

### 1.17 UPEC infection in pooled human urine

Apical-out hBOs withstand prolonged exposure to pooled human urine (PHU) with no significant increase in cell death over time compared with organoids maintained in standard L-WRN conditioned hBO medium (Figure 4A). Under these physiological conditions, the apical-out configuration enables direct access of uropathogenic bacteria to the epithelial surface without the need for microinjection.

**Figure 4:**
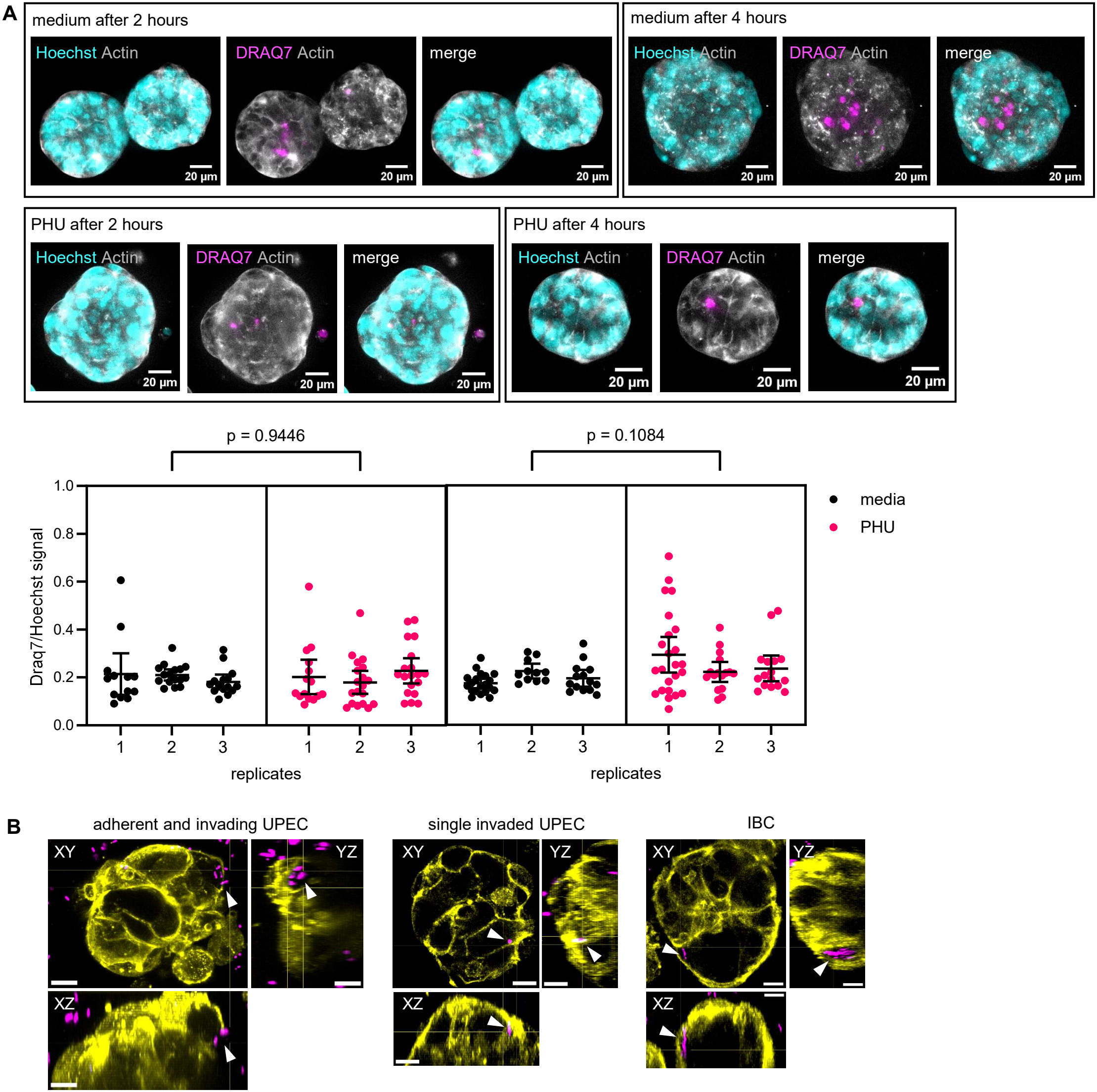
Apical-out human bladder organoids withstand pooled human urine exposure and can be infected with UPEC strain CFT073. A. Apical-out hBOs were sampled 2 hours and 4 hours post media change to either fresh medium (50/50 HBO/L-WRN) or pooled-human urine (PHU). Z-projection (maximum intensity) of representative images. Actin, grey; Hoechst, cyan; DRAQ7, magenta. DRAQ7 fluorescence was normalized to Hoechst fluorescence for each manually segmented organoid to account for differences in organoid size and cellular content. Nested t test (N = 3 biological replicates). Data shown as mean ± 95% confidence interval. Apical-out hBOs infected with UPEC strain CFT073. UPEC can adhere to and invade into apical-out hBOs, and form intracellular bacterial communities (IBCs) – highlighted with arrow heads. Actin, yellow; UPEC, magenta. Scale bar 10 µm.

Using the well-characterized UPEC strain CFT073 (Welch et al., 2002), the system recapitulates key infection phenotypes observed in vivo, including bacterial adherence, intracellular invasion and intracellular bacterial community (IBC)-like structures (Figure 4B and SI Figure 6).

### 1.18 Limitations

This protocol requires a rotating bioreactor system and compatible suspension culture tubes (e.g. CERO 3D system and CERO tubes, OLS). The current validation is restricted to H-6215 primary human bladder epithelial cells, which have previously been shown to robustly form physiologically relevant bladder microtissues in other systems (Paduthol et al., 2026; Tomasek et al., 2026).

Long-term culture beyond the described 8-day (+1 day after cryopreservation) maturation window has not been systematically evaluated, as the focus of this protocol is rapid and scalable generation of apical-out organoids. Furthermore, infection experiments were performed using the laboratory UPEC strain CFT073 only. Thus, the applicability of the system to additional clinical UPEC isolates and uropathogens remains to be determined.

### 1.19 Troubleshooting

For a list of potential problems and their respective solutions, see Table 6.

**Table 6.** Potential problems, reasons and solutions.

| Problem | Possible reason | Solution |
|---|---|---|
| excess organoid aggregation/clumping | insufficient dispersion of Matrigel droplets (day 2 and day 6/thaw day+1) | pipette 300-500 µl repeatedly (20x) |
| organoids stick to plastic | no BSA coating | pre-coat laboratory plastics |
| poor differentiation | passage number too high | use P6-P8 at maximum |
|  | incorrect media preparation | use a 50/50 ratio of HBO media and L-WRN conditioned medium |
| slow growth or abnormal development | mycoplasma contamination | test routinely for mycoplasma contamination |
| organoid fusion/non-circular shaped organoids | incorrect rotation | verify CERO 3D settings |
|  | insufficient spontaneous droplet formation of Matrigel | use warm cell-containing HBO medium and ice-cold Matrigel-containing L-WRN conditioned medium |
|  | wrong Matrigel concentration | use 4% Matrigel for 5 days, then decrease to 2% |
| low yield | insufficient cell number at seeding | seed 1 million cells per tube |
|  | organoids loss due to adhesion to plastic | coat laboratory plastics with BSA |
| no infection phenotype observed | residual antibiotics (e.g. Primocin) interfering with bacterial growth | remove residual Primocin prior to infection with bacteria |

## Discussion

Here, we describe a rapid and scalable protocol for the generation of apical-out human bladder organoids (hBOs) from primary epithelial cells in a rotating bioreactor, requiring only minimal extracellular matrix support and operator handling. Within eight days, cells seeded in suspension self-organize into solid, stratified organoids that display an outward-facing umbrella-like cell layer, thereby exposing the apical surface that is otherwise sequestered within the central lumen of conventional dome-grown organoids. This inversion of polarity, together with the speed and simplicity of the workflow, addresses within a single system two long-standing constraints of urothelial organoid models – restricted apical accessibility and limited scalability.

A defining feature of this model is the degree to which it recapitulates the stratification and terminal differentiation of the native bladder epithelium. Combining RT-qPCR, whole-mount and section-based immunofluorescence, histology and transmission electron microscopy, we find that mature hBOs reproduce the spatial organization of the principal urothelial compartments: terminally differentiated markers (UP3A, CK20) are preferentially confined to the outer, apical-facing umbrella-like layer, the intermediate marker CK13 occupies the suprabasal layers and the basal marker CK5 is found towards the organoid inside. Junctional localization of ZO-1 is consistent with the establishment of epithelial barrier structures, while electron microscopy confirms a compact, multilayered architecture without a central lumen or necrotic core, while rather indicating the presence of actively dividing cells. Importantly, this organization was reproducible across independent operators, and it was preserved in organoids that had been cryopreserved and subsequently recovered (D8+1), indicating that differentiation fidelity is a robust property of the system rather than an operator- or batch-dependent outcome.

Beyond fidelity, this protocol for suspension-grown organoids formed in rotation culture offers substantial practical advantages over conventional Matrigel-dome cultures (Tomasek et al., 2026) including reduced culture time, increased scalability, and decreased requirements for extracellular matrix and media (Table 7): A single CERO tube yields approximately 216,000 organoids – roughly 1.8-fold more per culture than a 24-well dome plate – while reducing media consumption around six-fold and Matrigel usage approximately 1.7-fold, together with shortening the time to maturation from roughly two weeks to eight days. Equally relevant for routine use, the suspension format largely eliminates the delicate, dome-by-dome handling that conventional cultures require, cutting operator hands-on time for organoid establishment. However, most notably, the apical-out configuration removes the requirement of technically challenging and low-throughput microinjections to access the apical surface. The option to cryopreserve the bulk of a culture at day 5 and recover a defined fraction on demand further decouples organoid production from experimental scheduling.

**Table 7.**
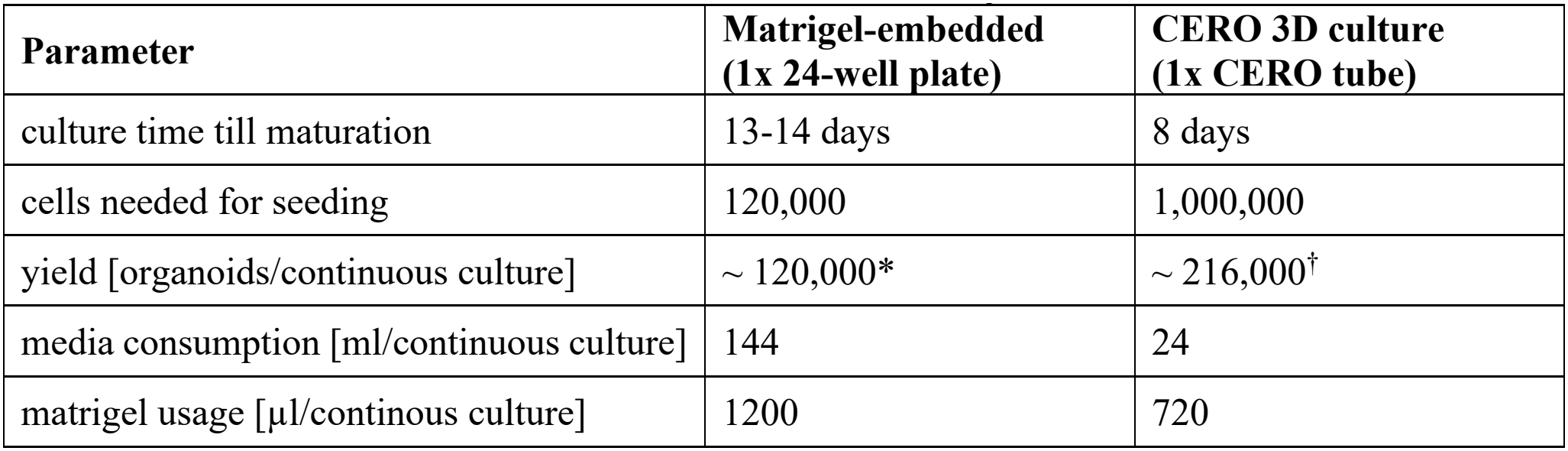
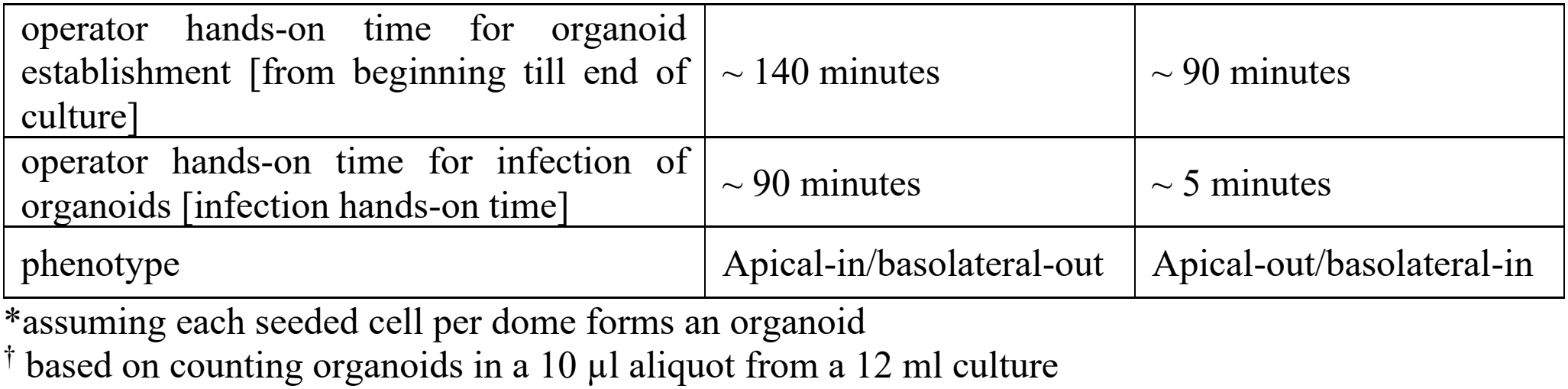
Parameters of conventional dome culture hBOs and suspension ones.

The apical-out configuration is particularly advantageous for infection biology and drug treatment assays. Because the umbrella-like cell layer faces the medium, uropathogenic bacteria gain direct access to the physiologically relevant apical surface without the technically demanding microinjection required in apical-in dome organoids. We show that hBOs tolerate prolonged exposure to 100% pooled human urine at intermediate osmolarity without a significant increase in cell death, allowing infection to be studied under near physiological conditions. Under these conditions, infection with the well-characterized UPEC strain CFT073 reproduced key hallmarks of bladder infection, including bacterial adherence, intracellular invasion and the formation of intracellular bacterial community-like structures. The model is therefore compatible with standard readouts such as gentamicin protection assays and high-resolution imaging, and it lowers both the equipment and skill barriers to performing physiologically contextualized infection assays on human primary urothelium.

Several limitations should nonetheless be considered when adopting this protocol. The method depends on a rotating bioreactor and compatible suspension tubes, and our validation is currently restricted to one commercial source of primary human bladder epithelial cells and one laboratory-adapted UPEC isolate; extension to additional UPEC isolates or other uropathogens, as well as patient-derived bladder cells or induced pluripotent stem cell-derived urothelium will be needed to establish generality. Our protocol was deliberately optimized for rapid maturation and has not been evaluated for long-term maintenance beyond the eight-day window (plus one day after cryorecovery), and the fold-changes in differentiation markers, while directionally consistent across assays, varied between biological replicates although did not reach statistical significance – a variability inherent to primary cells that users should anticipate.

Looking forward, we anticipate that this platform will be broadly useful wherever direct and reproducible access to differentiated human urothelium is required. The open apical surface makes the system well suited to screening applications, such as for antimicrobials, host-directed therapies or barrier-modulating compounds, and to studies of apical membrane biology that are difficult to address in enclosed-lumen models. More broadly, by combining native-like stratification with apical accessibility, scalability and a markedly simplified workflow, apical-out hBOs provide a practical and adaptable platform for dissecting bladder epithelial biology and host-pathogen interactions, and for bridging mechanistic organoid studies with the physiological context of the human urinary tract.

## Author Contributions

Conceptualization: VJV, KT. Data curation: VB, VJV, HS, KT. Formal analysis: VB, VJV, HS, KT. Funding acquisition: JDM. Investigation: VB, VJV, HS, KT. Methodology: VB, VJV, KT. Project administration: KT. Resources: LL, JDM. Supervision: KT. Validation: VB, VJV, KT. Visualization: VJV, HS, KT. Writing original draft: VB, KT. Writing review & editing: all authors

## Acknowledgments

We thank Dr. Andrea Krapp from the SCOL at EPFL for scientific support and the EPFL MultiOmics Histology Core Facility, Damien De Bellis at UNIL for preparing TEM sections and the UNIL Electron Microscopy Facility, as well as Joris Veger from UZH, and Hatice Demiral and Dr. Markus Uhring from Omni Life Sciences for their technical support. We thank all members of the McKinney laboratory for scientific discussions.

## Supplementary Material

The Supplementary Material for this article is provided as a separate file and includes Supplementary Figures SI 1-6.

## Data Availability Statement

All datasets linked to this study will be made available upon acceptance. Raw and processed microscopy imaging data will be made available at the BioImage Archive. Quantified results and statistical analyses will be made available at Zenodo. Analysis codes will be made available at Zenodo and GitHub.

## Declaration of generative AI and AI-assisted technologies in the manuscript preparation process

During the preparation of this work the author(s) used Claude (Anthropic) in order to assist with language refinement and clarity in sections of the manuscript, including improving phrasing, grammar and flow. AI was also used for the code generation for image analysis. After using this tool/service, the author(s) reviewed and edited the content as needed and take(s) full responsibility for the content of the published article.

