## Supplementary Figures for "A simplified protocol for the generation of apical-out human bladder organoids in suspension using a rotating bioreactor"

**SI Figure 1**

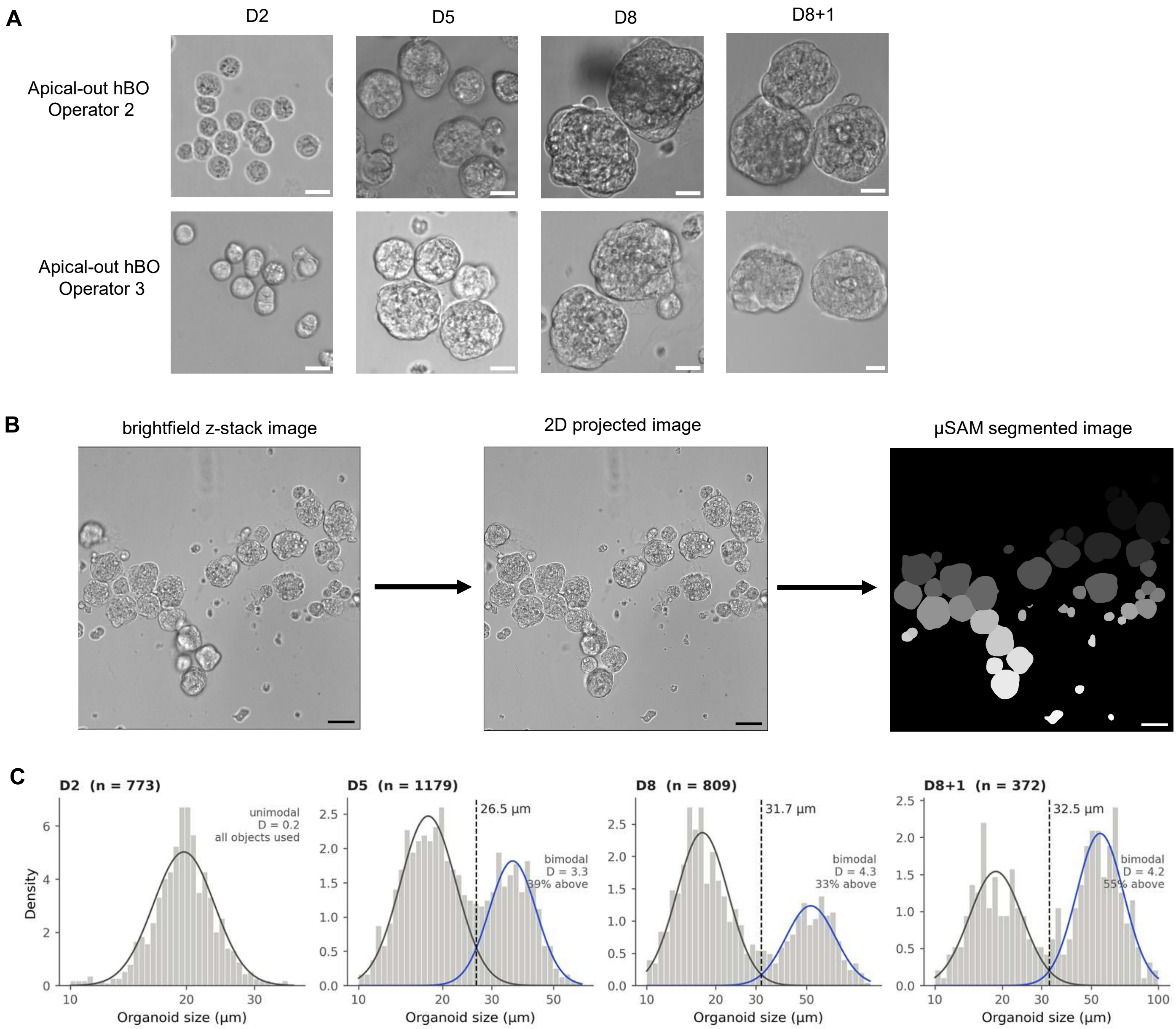
A
D2
D5
D8
D8+1
Apical-out hBO Operator 2
Apical-out hBO Operator 3
B
brightfield z-stack image
2D projected image
µSAM segmented image
C
D2 (n = 773)
unimodal
D = 0.2
all objects used
Density
Organoid size (µm)
D5 (n = 1179)
26.5 µm
bimodal
D = 3.3
39% above
D8 (n = 809)
31.7 µm
bimodal
D = 4.3
33% above
D8+1 (n = 372)
32.5 µm
bimodal
D = 4.2
55% above

SI Figure 1: Organoid brightfield image analysis.

A. Organoid brightfield image replicates from two additional operators across growth timeline starting from Day 2 in culture (D2), progressing through D5, D8 grown continuously, and D8+1 following freeze-thaw protocol, i.e. frozen on Day 5 and cultured for 3 additional days after thawing. Scale bar 20 µm.
B. Brightfield z-stack, projected into a single 2D image with all organoids in focus and segmented using µSAM. Organoids touching image borders are ignored, and masks are manually curated to disregard poorly segmented organoids. Scale bar 50 µm.
C. Rationale for unimodal versus bimodal distribution of organoid sizes based on fitting one versus two Gaussian curves on the organoid size distribution.

**SI Figure 2**

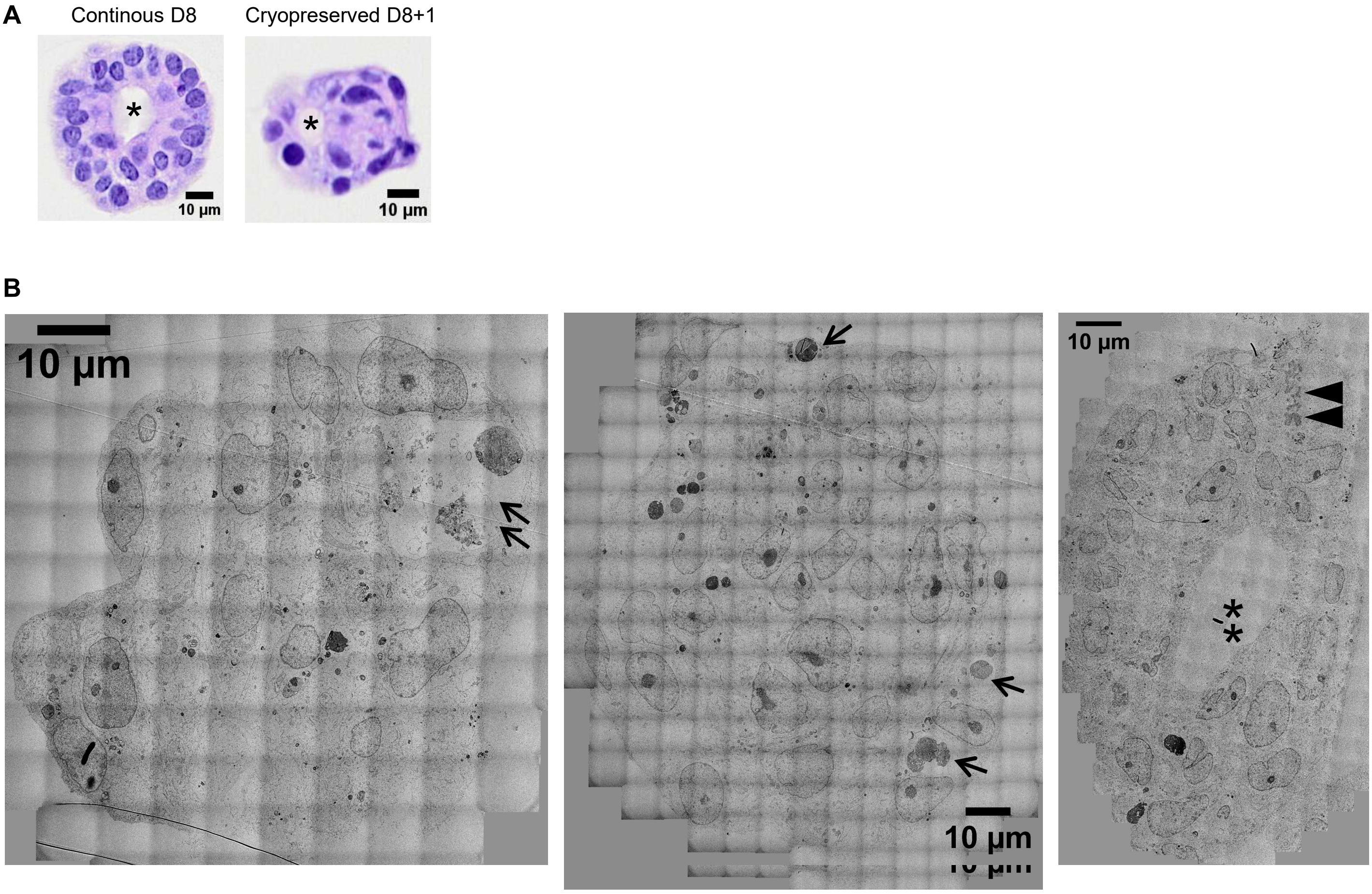

SI Figure 2: Replicate H&E images of organoids.

A. H&E (Hematoxylin & Eosin) images of continuous (D8) and cryopreserved (D8+1) organoids. A small fraction of organoids incorporate larger aggregates of Matrigel (indicated with an asterisk), while cells start growing around them during development.
B. Transmission electron microscopy images of continuously grown (D9) apical-out hBOs. Some putatively apoptotic cells are indicated by arrows, while the arrowhead points to a mitotic event in a dividing cell. The asterisk shows a BME aggregate.

**SI Figure 3**

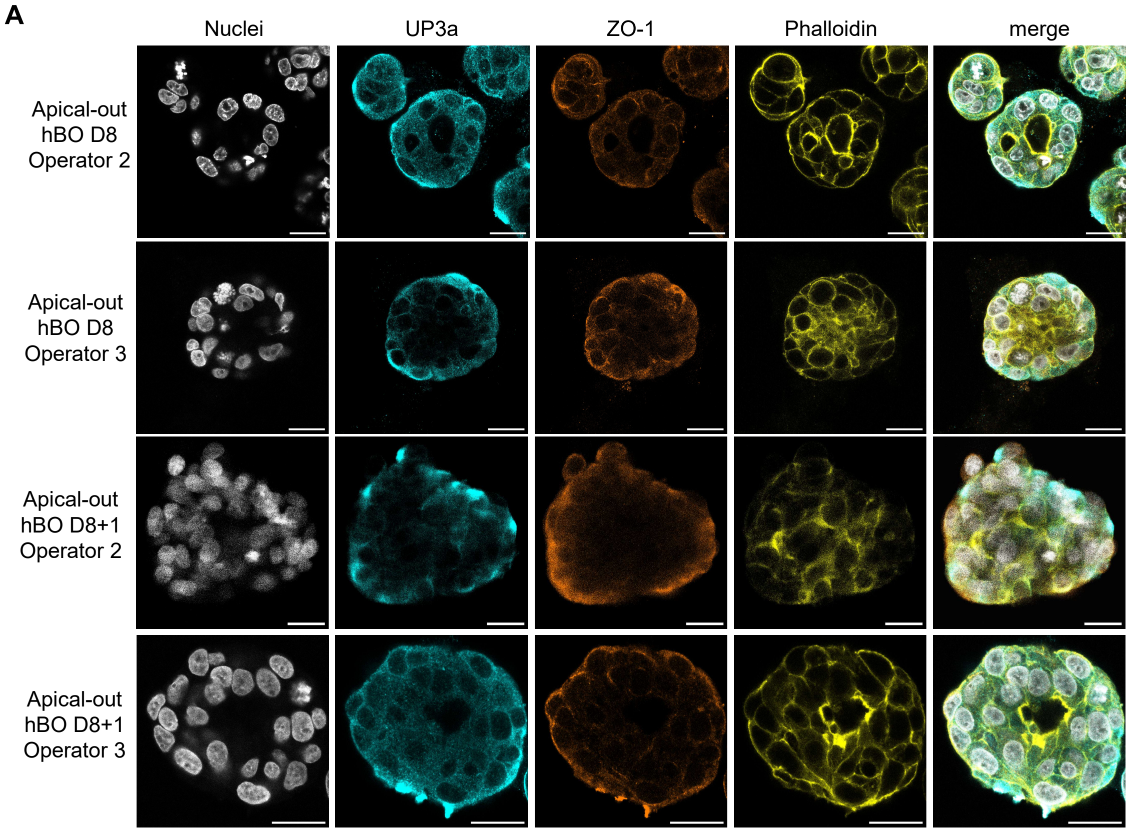
A
Nuclei
UP3a
ZO-1
Phalloidin
merge
Apical-out hBO D8 Operator 2
Apical-out hBO D8 Operator 3
Apical-out hBO D8+1 Operator 2
Apical-out hBO D8+1 Operator 3

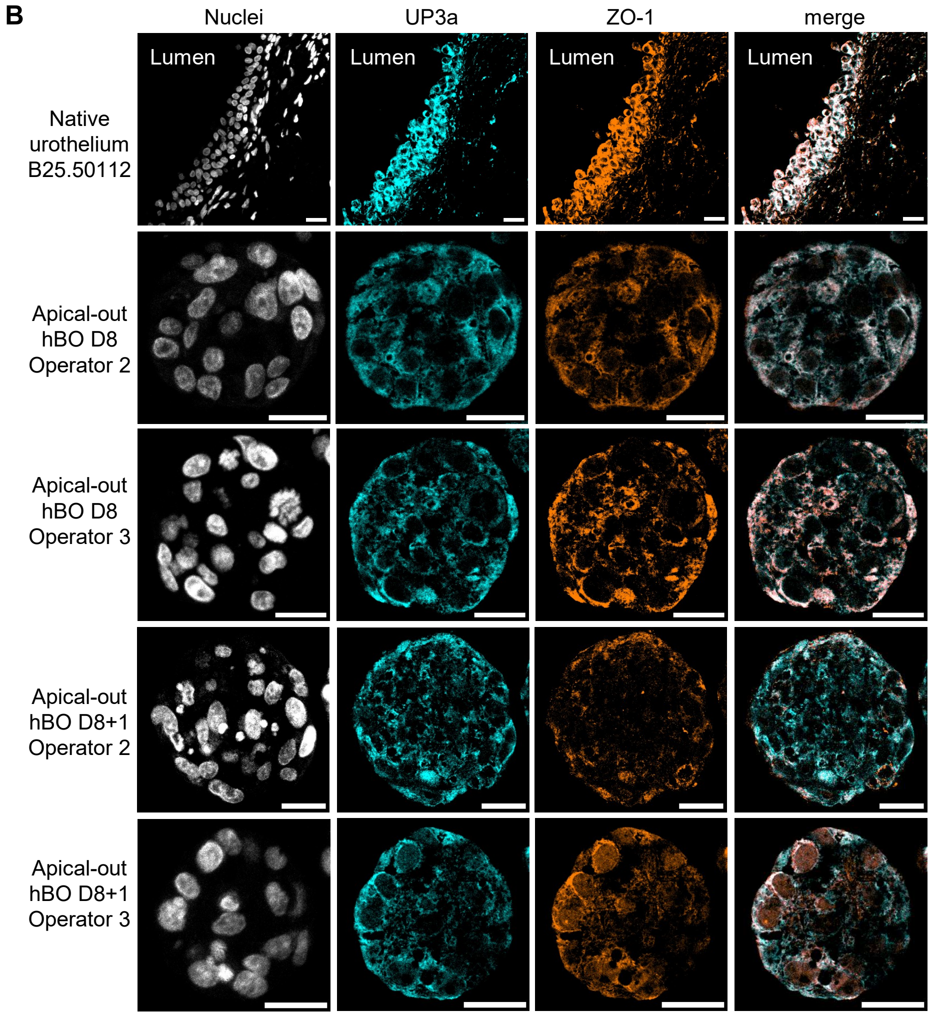
B
Nuclei
UP3a
ZO-1
merge
Lumen
Lumen
Lumen
Lumen
Native urothelium B25.50112
Apical-out hBO D8 Operator 2
Apical-out hBO D8 Operator 3
Apical-out hBO D8+1 Operator 2
Apical-out hBO D8+1 Operator 3

SI Figure 3: Replicate images of organoids stained for UP3a and ZO-1.

Immunofluorescent images of organoids. Representative images of (A) whole mount processed apical-out human bladder organoids and (B) 4 μm sections of native urothelium and apical-out human bladder organoids across different operators. Nuclei, grey. UP3a, cyan. ZO-1, amber. Scale bar 20 μm.

### SI Figure 4

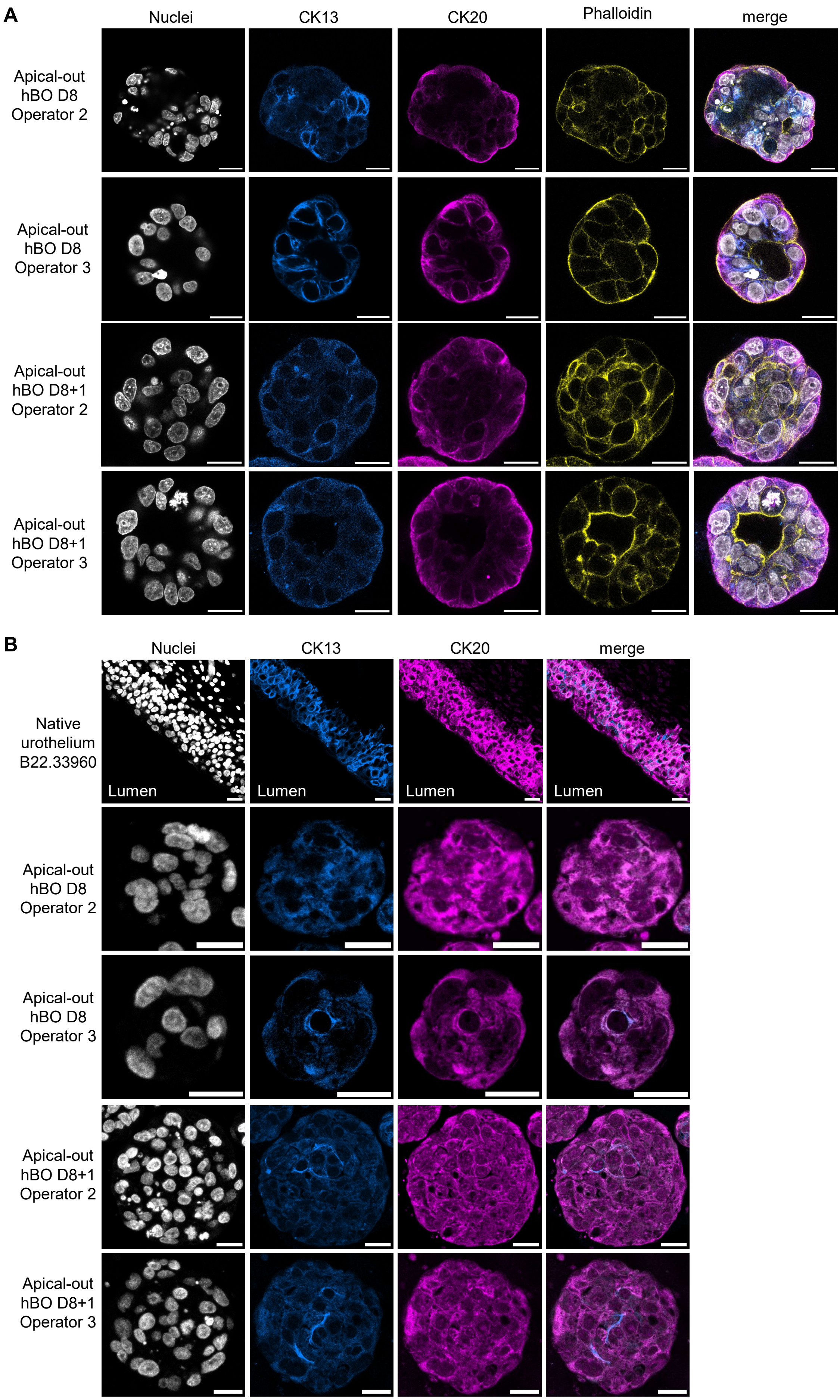
A
Nuclei
CK13
CK20
Phalloidin
merge
Apical-out hBO D8 Operator 2
Apical-out hBO D8 Operator 3
Apical-out hBO D8+1 Operator 2
Apical-out hBO D8+1 Operator 3
B
Nuclei
CK13
CK20
merge
Native urothelium B22.33960
Lumen
Lumen
Lumen
Lumen
Apical-out hBO D8 Operator 2
Apical-out hBO D8 Operator 3
Apical-out hBO D8+1 Operator 2
Apical-out hBO D8+1 Operator 3

SI Figure 4: Replicate images of organoids stained for CK20 and CK13.

Immunofluorescent images of histology sections. Representative images of (A) whole mount processed apical-out human bladder organoids and (B) 4 µm sections of native urothelium and apical-out human bladder organoids across different operators. Nuclei, grey. CK13, azure. CK20, magenta. Scale bar 20 µm.

**SI Figure 5**

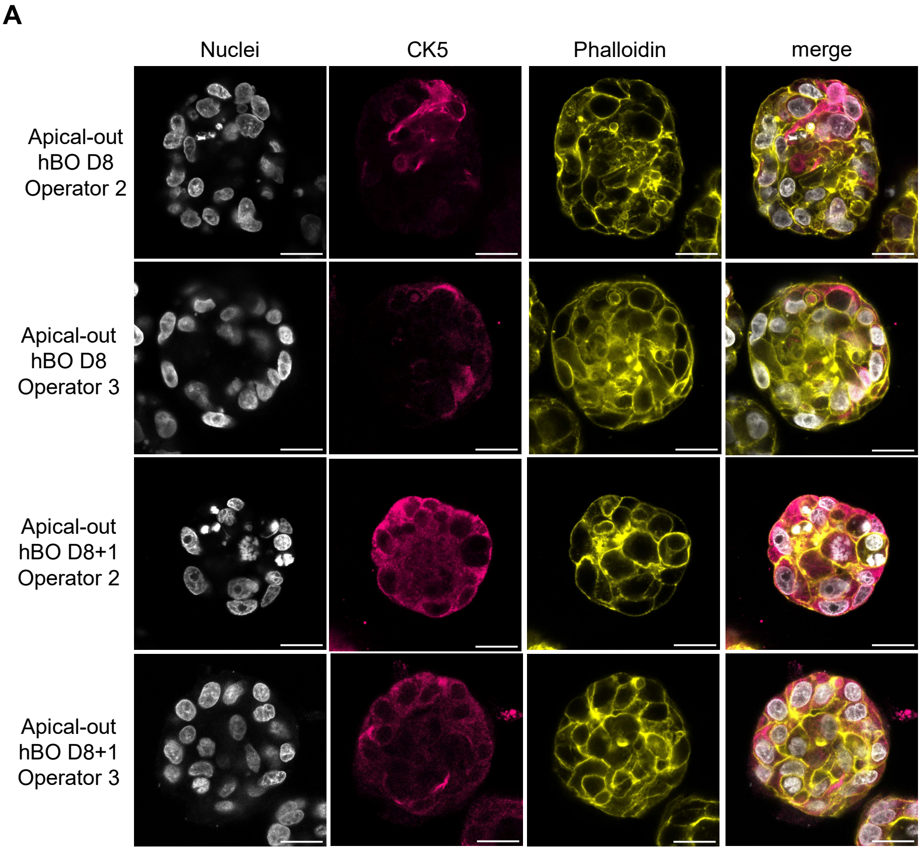
A
Nuclei
CK5
Phalloidin
merge
Apical-out hBO D8 Operator 2
Apical-out hBO D8 Operator 3
Apical-out hBO D8+1 Operator 2
Apical-out hBO D8+1 Operator 3

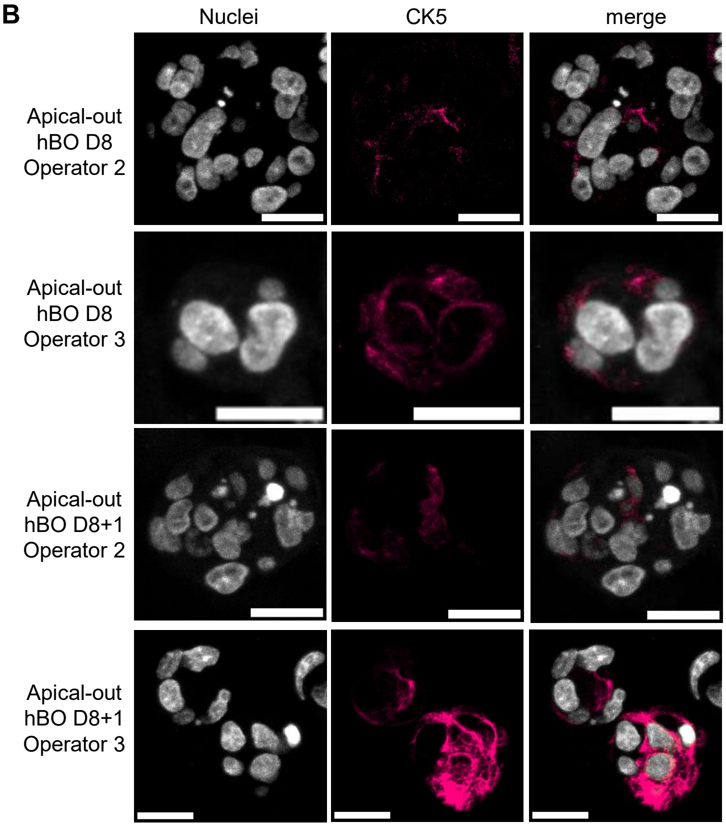
B
Nuclei
CK5
merge
Apical-out hBO D8 Operator 2
Apical-out hBO D8 Operator 3
Apical-out hBO D8+1 Operator 2
Apical-out hBO D8+1 Operator 3

SI Figure 5: Replicate images of organoids stained for CK5.

Immunofluorescent images of histology sections. Representative images of (A) whole mount processed and (B) 4 μm sections of apical-out human bladder organoids across different operators. Nuclei, grey. CK5, pink. Scale bar 20 μm.

**SI Figure 6**

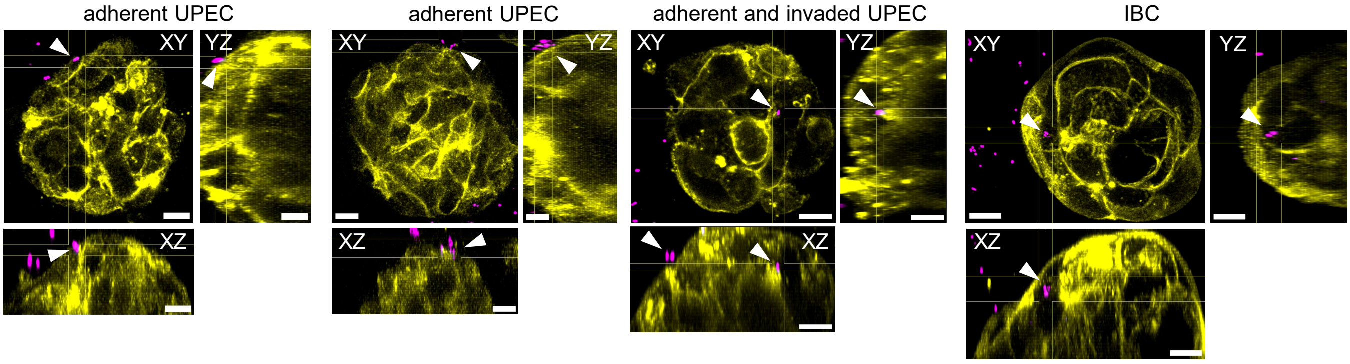

adherent UPEC
adherent UPEC
adherent and invaded UPEC
IBC
XY
YZ
XZ
XY
YZ
XZ
XY
YZ
XZ
XY
YZ
XZ

SI Figure 6: Replicate images of infected organoids.

Replicate images of apical-out hBOs infected with UPEC strain CFT073. UPEC can adhere to and invade into apical-out hBOs, and form intracellular bacterial communities (IBCs) – highlighted with arrow heads. Actin, yellow; UPEC, magenta. Scale bar 10 µm.
